# Effects of Aging on Cortical Representations of Continuous Speech

**DOI:** 10.1101/2022.08.22.504825

**Authors:** I.M Dushyanthi Karunathilake, Jason L. Dunlap, Janani Perera, Alessandro Presacco, Lien Decruy, Samira Anderson, Stefanie E. Kuchinsky, Jonathan Z. Simon

## Abstract

Understanding speech in a noisy environment is crucial in day-to-day interactions, and yet becomes more challenging with age, even for healthy aging. Age-related changes in the neural mechanisms that enable speech-in-noise listening have been investigated previously; however, the extent to which age affects the timing and fidelity of encoding of target and interfering speech streams are not well understood. Using magnetoencephalography (MEG), we investigated how continuous speech is represented in auditory cortex in the presence of interfering speech, in younger and older adults. Cortical representations were obtained from neural responses that time-locked to the speech envelopes using speech envelope reconstruction and temporal response functions (TRFs). TRFs showed three prominent peaks corresponding to auditory cortical processing stages: early (∼50 ms), middle (∼100 ms) and late (∼200 ms). Older adults showed exaggerated speech envelope representations compared to younger adults. Temporal analysis revealed both that the age-related exaggeration starts as early as ∼50 ms, and that older adults needed a substantially longer integration time window to achieve their better reconstruction of the speech envelope. As expected, with increased speech masking, envelope reconstruction for the attended talker decreased and all three TRF peaks were delayed, with aging contributing additionally to the reduction. Interestingly, for older adults the late peak was delayed, suggesting that this late peak may receive contributions from multiple sources. Together these results suggest that there are several mechanisms at play compensating for age-related temporal processing deficits at several stages, but which are not able to fully reestablish unimpaired speech perception.

**NEW & NOTEWORTHY:** We observed age-related changes in cortical temporal processing of continuous speech that may be related to older adults’ difficulty understanding speech in noise. These changes occur in both timing and strength of the speech representations at different cortical processing stages, and depend on both noise condition and selective attention. Critically, their dependency on noise condition changes dramatically among the early, middle, and late cortical processing stages, underscoring how aging differentially affects these stages.

## INTRODUCTION

Speech communication is crucial in day-to-day interactions, and our interactions with others depend heavily on our ability to understand speech in a variety of listening conditions. Speech comprehension becomes increasingly difficult in a noisy environment, and, critically, degrades further with aging. Compared to younger adults, older adults have been observed to exhibit greater difficulty in auditory tasks in the presence of background noise, whether in relatively simple paradigms such as pitch discrimination (Fitzgibbons and Gordon-Salant 1995) and gap detection (Snell 1997), or in acoustically complex paradigms such as speech listening in noise (Frisina and Frisina 1997; Gordon-Salant et al. 2006). Because poor speech comprehension in noise is associated with adverse psycho-social effects (Bess et al. 1989; Strawbridge et al. 2000), depression (Gopinath et al. 2009) and dementia (Uhlmann et al. 1989), identifying age-related changes in the neural mechanisms that underlie speech-in-noise difficulties may be critical for developing remediations that improve communication and quality of life among older adults. The present study aims to investigate the timing and fidelity of cortical auditory processing in more ecologically valid naturalistic speech paradigm that could be incorporated into both diagnostic evaluations and treatments aimed at speech comprehension problems in older adults.

Numerous studies have demonstrated important age-related anatomical and functional changes in auditory cortical pathways that may contribute to listening difficulties. In studies with human participants, exaggerated neural activity in the auditory cortex has been observed in older adults during speech and speech-in-noise tasks (Presacco et al. 2016a; Manan et al. 2017; Brodbeck et al. 2018; Decruy et al. 2019; Mesik et al. 2021; Gillis et al. 2023), even when age-related hearing loss is mild, potentially reflecting changes in the central auditory system that could contribute to speech comprehension difficulties in older adults. Animal studies have also reported age-related excitatory and inhibitory imbalance in auditory cortex (Hughes et al. 2010a; Caspary et al. 2013), leading to altered neural coding.

Age-related changes in auditory, linguistic, and cognitive processes can all influence speech understanding (for a review, see Kuchinsky and Vaden (2020)). Age-related changes in cognitive functions limit speech understanding among older adults (Dryden et al. 2017), in addition to other combinations of anatomical, functional and cognitive factors (Schneider and Pichora-Fuller 2000). Therefore, audiometric measurements of the peripheral auditory system alone may not be sufficient to evaluate and manage the speech understanding difficulties reported by older adults.

The human neurophysiology underlying age-related changes in the timing and fidelity of encoding of connected speech is not well understood. High temporal resolution recording methods such as magnetoencephalography (MEG) and electroencephalography (EEG) are well-suited to accurately estimate such neural responses and the effects of aging on them. Much of this research routinely focuses on auditory evoked responses that quantify the central auditory functions obtained by averaging response to many repetitions of simple stimuli. Such studies often find that the cortical auditory event potential (CAEP), e.g., the P1-N1-P2 complex, is enhanced with aging (Tremblay et al. 2002, 2003; McCullagh and Shinn 2013; Bidelman et al. 2014). Reduced inhibitory neurotransmitters in the ascending auditory pathway may contribute to these enhanced neural response (Tremblay and Ross 2007). In addition, these studies have also reported delayed N1 and P2 peaks in older adults compared to younger adults (for a review, see Anderson and Karawani (2020)). Enhanced and delayed neural responses may demonstrate at least partial compensation for and restoration of degraded subcortical input at the cortex.

However, simple stimuli (e.g., tones, clicks, and single speech syllables) do not well replicate real-world listening where the ultimate goal is speech comprehension (Keidser et al. 2020). Computational tools have been developed, however, that can analyze neural responses to *continuous speech*, typically in terms of speech encoding and decoding models. Speech envelope reconstruction analysis and temporal response function (TRF) analysis respectively measure neural speech processing as linear decoding and encoding methods, and can be used for both attended (foreground) and unattended (background) speech streams. In contrast CAEP responses cannot straightforwardly segregate the neural responses to individual acoustic components in the stimulus (e.g., latency effects of selective attention). In this study, we extend the limitations of CAEP studies using more naturalistic speech paradigm and by investigating the timing and fidelity of both attended and unattended speech responses that could explain cortical processing differences in older adults.

In recent years, both EEG and MEG studies analyzing the reconstruction accuracy of the speech envelope have reported an enhanced cortical representation of the attended speech envelope in older adults. Reconstruction analysis integrates over a long time window (typically 500 ms), however, making estimating the latency time-course of age-related over-representation more difficult. Presacco et al. (2016a) showed that older adults’ cortical ability to track the speech envelope is significantly reduced when decreasing the integration window to 150 ms, indicating that, at least, longer latencies contribute significantly. In this study, we replicate and extend the temporal analysis window results of Presacco et al. (2016a) using a nonlinear mixed effects modeling approach (i.e., generalized additive mixed effects models, GAMMs) to better understand the time course of speech processing.

While temporal integration window analysis is robust because of its integration over sensors and time, detailed temporal information can be gained directly from a TRF analysis (Ding and Simon 2012a; Power et al. 2012). Prominent peaks in the TRF, the M50_TRF_, M100_TRF_ and M200_TRF_, can be ascribed to different auditory processing stages in the cortex with the corresponding latencies (Lister et al. 2011). In analogy to the P1-N1-P2 CAEP peaks at the corresponding latencies, it has been suggested that the early M50_TRF_ (P1-like) peak dominantly reflects the low-level acoustic features such as changes in acoustic power (Näätänen and Winkler 1999; Ceponiene et al. 2005), whereas the M100_TRF_ (N1-like) peak reflects processing of selectively attended features (Näätänen and Winkler 1999). Similarly, the M50_TRF_ has been shown to depend more on the properties of the acoustic stimulus than the focus of selective attention, whereas the M100_TRF_ shows the opposite (Ding and Simon 2012b, 2013). The late peak M200_TRF_ (P2-like) may reflect stimulus familiarity (Sheehan et al. 2005) and training (Tong et al. 2009); it is quite late for encoding bottom-up acoustic features, but appropriately positioned to reflect a representation of auditory object formation (Näätänen and Winkler 1999). In this study, we investigate how amplitudes and latencies of each of these processing stages are affected by aging and listening condition in a continuous speech paradigm, to better understand the age-related temporal processing differences.

In comparison to the neural representations of the attended speech envelope discussed above, neural representations of unattended speech in older listeners are less well understood (though see Zan et al. (2020)). In this study, we extend the timing and fidelity analysis to unattended speech to understand how aging affects auditory scene representation in general, and stream segregation in particular, in the cortex.

In summary, this study aims to systematically investigate age-related neurophysiological effects on continuous speech processing, using both envelope reconstruction and TRF analysis. The effects of age, selective attention, and competing speech masking are evaluated concurrently. To minimize the effect of age-related peripheral hearing loss, only participants who had hearing thresholds within clinically normal limits through 4.0 kHz were recruited in the study. Expanding beyond prior results, we expected to observe exaggerated representation of the speech envelope in older adults irrespective of the competing speech, selective attention and processing stage. As older adults show difficulty understanding speech-in-noise, we expect that timing and fidelity measures will be differently or severely affected from speech in quiet to speech in noise, in older adults compared to younger adults. Finally, for the late neural processing stages, we expect that older adults’ neural measures patterns will grow from younger adults. (e.g., enhanced top-down connectivity associated with cognitive compensation in older adults).

## METHODS

All experiments were performed in accordance with the guidelines and regulations for human subject testing by the University of Maryland’s Institutional Review Board. All participants gave written informed consent to participate in the protocol and they were compensated for their time.

### Participants

34 native English speakers participated in the study: 18 younger adults (7 males; mean age 20 y, range 18-26 y) and 16 older adults (5 males; mean age 70 y, range 65-78 y). Data from two additional subjects (1 older and 1 younger) were not included in the analysis due to data saturation caused by excessive MEG artifacts. All participants had normal hearing (see Figure 1), defined as pure-tone thresholds ≤ 25 dB hearing level (HL) from 125 to 4000 Hz in at least one ear, and no more than 10 dB difference between ears at each frequency. Only subjects with Montreal Cognitive Assessments (MoCA) scores within normal limits (≥ 26) and no history of neurological disorder were included.

**Figure 1.**
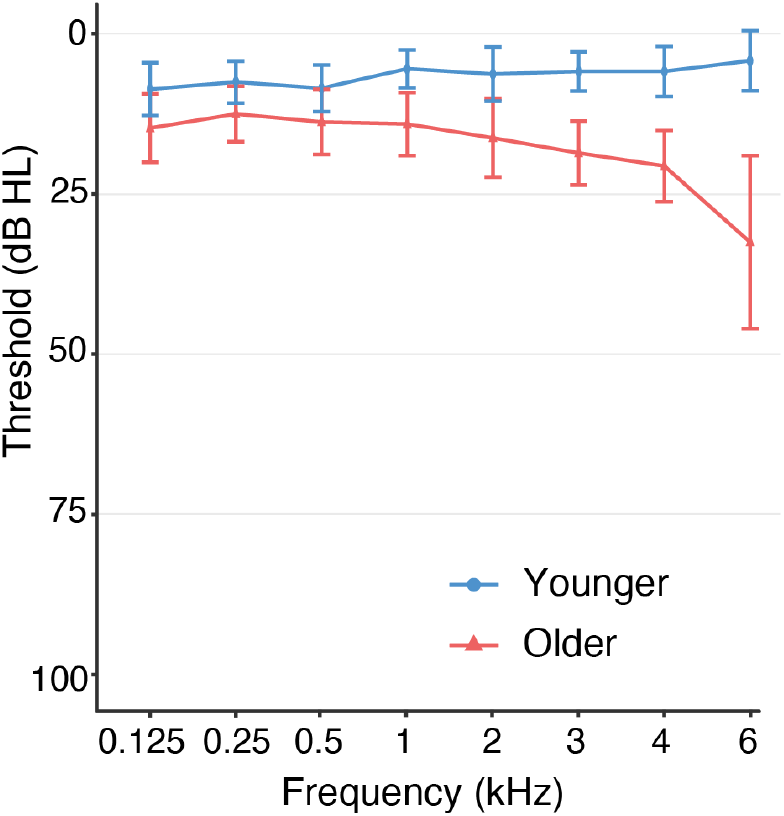
Audiogram of the grand average across ears for younger (blue) and older (red) participants. Error bars indicate ± one standard deviation. Both groups have clinically normal hearing, defined as pure-tone thresholds ≤ 25 dB HL from 125 - 4000 Hz in at least one ear, and no more than 10 dB difference between ears at each frequency.

### Stimuli and experimental design

The speech segments were extracted from the audio book, “The Legend of Sleepy Hollow”, by Washington Irving, narrated by separate male (https://librivox.org/the-legend-of-sleepy-hollow-by-washington-irving) and female (https://www.amazon.com/The-Legend-of-Sleepy-Hollow-audiobook/dp/B00113CMHE) talkers. Talker pauses greater than 400 ms were shortened to 400 ms and then recordings were low pass filtered <4 kHz using a third order elliptic filter. One-minute-long audio stimulus segments (22.05 kHz sampling rate) were extracted and rms value of the sound amplitudes was normalized to have equal perceptual loudness in MATLAB (MathWorks) (Ding and Simon 2012b). All stimuli were presented diotically (identically in each ear).

Four types of stimuli were presented: single talker (“quiet speech”), two talkers (“competing talkers”) at two different relative loudness levels (0 dB SNR and −6 dB SNR), and single talker mixed with three-talker babble (“babble speech”). For the competing talker speech trials (a trial was defined as a 60 s duration speech passage presentation), participants were asked to selectively attend to one talker while ignoring the other, for which there were two signal-to-noise ratio (SNR) levels 0 dB and −6 dB. For the babble condition, only the female talker was used as foreground, with the three-talker babble mixed in at 0 dB SNR. In the mixed speech and babble speech conditions, the sound level of the attended talker was identical to that of the corresponding single talker condition; only the sound level for the unattended talker or babble was altered to change the noise condition. The order of the four competing-talker blocks was counterbalanced across subjects in the order of attended-talker and SNR (2 × 2). The babble speech condition was always presented as the third block. In the competing talker and babble speech conditions, each stimulus was presented three times. At the end of each of these blocks, the attended and unattended speech stimuli in that block were presented alone as single talker speech without repetition (for babble speech condition, only the attended female talker speech was presented); otherwise, no speech segment was ever re-used across blocks.

Sound level was calibrated to approximately 70 dBA sound pressure level (SPL) using 500 Hz tones and equalized to be approximately flat from 40 Hz to 4 kHz. The stimuli were delivered with E-A-RTONE 3 A tubes (impedance 50 Ω), which strongly attenuate frequencies above 3-4 kHz, and E-A-RLINK (Etymotic Research, Elk Grove Village, United States) disposable earbuds inserted into the ear canals.

To motivate the participants to engage in the task, at the end of each trial, a simple story-content question based on the attended passage was asked. After the first trial of each condition, participants were also asked to rate the intelligibility rating on a scale from 0 to 10 (0 being completely unintelligible and 10 being completely intelligible). This estimated rating was used as a subjective measure of intelligibility.

### Data recording

Non-invasive neuromagnetic responses were recorded using a 157 axial gradiometer whole head MEG system (KIT, Kanazawa, Japan), inside a dimly lit, magnetically shielded room (Vacuumschmelze GmbH & Co. KG, Hanau, Germany) at the Maryland Neuroimaging Center. The data were sampled at 2 kHz along with an online anti-aliasing low-pass filter with cut off frequency at 500 Hz and a 60 Hz notch filter. Three separate channels function as environmental reference channels. Subjects lay supine during the entire experiment and were asked to minimize body movements. During the task, subjects were asked to keep their eyes open and fixate on a male/female cartoon face at the center of screen, corresponding to the attended talker. Pupil size data were recorded simultaneously with an eye tracker (EyeLink 1000 Plus); those results will be presented separately.

### Data processing

All data analysis was conducted in MATLAB R2020a. The raw MEG data was first denoised by removing non-functioning and saturated channels, and then with Time Shifted Principle Component Analysis (TSPCA) (de Cheveigné and Simon 2007), using the three reference channels to project out environmental magnetic noise not related to the brain activity, and then with Sensor Noise Suppression (SNS) (de Cheveigné and Simon 2008a) to project out sensor specific noise. To focus on low frequency cortical activity, the remaining data was band pass filtered between 1-10 Hz with an order 6000 Hamming-windowed finite impulse response filter (FIR), and compensated for group delay. A blind source separation method, Denoising Source Separation (DSS) (Sarela and Valpola 2005; de Cheveigné and Simon 2008) was next applied to the repeated trials to extract those subject-specific spatial components that are reliable over trials, ranked in order of reproducibility. The first six DSS components were used for the stimulus reconstruction analysis. Generally, the first component corresponds to the primary auditory component (i.e., with a topography consistent with bilateral temporal lobe sources) and so was selected for all subsequent TRF estimation; for one older adult, the second component reflected the primary auditory component and so was selected for TRF estimation in place of the first. As the polarity of a raw DSS component is arbitrary, which might affect TRF peak estimation and comparison between subjects, we aligned the polarity of the primary auditory component across subjects by flipping both the sign and the topography of that component for any subject whose topography gave a negative spatial correlation with a fiducial topography (i.e., the topography produced by an evoked response to a pure tone with standard M50/M100 peaks). Finally, data were downsampled to 250 Hz for TRF analysis and to 100 Hz for the stimulus reconstruction analysis.

The envelope of the audio waveform was processed to match the processed MEG data. Each attended and unattended single talker stimulus was downsampled to 2 kHz and the logarithmic envelope was extracted as described in Biesmans et al. (2017). Then the envelope was filtered with the same bandpass filter (1–10 Hz) applied to the MEG data (and group delay compensation) and downsampled to 250 Hz and 100 Hz for TRF or stimulus reconstruction analysis, respectively.

### Behavioral tests

#### Flanker Test

The ability to selectively attend to one talker and ignore (inhibit) the other requires executive function. The Flanker Inhibitory Control and Attention Test of the National Institute of Health Toolbox (Gershon et al. 2013) was used as a measure of the subject’s general behavioral ability to suppress competing stimuli in a visual scene. Participants were instructed to identify the direction of a central arrow while ignoring the directions of a surrounding set of four arrows (“flankers”) by pressing a key as quickly and accurately as possible. The direction of the central arrow could be similar (congruent) or different (incongruent) to the surrounding arrows. The unadjusted scale score was calculated based on the reaction times (RTs) and the accuracy. Higher flanker scores represent better performance.

#### Speech-In-Noise (SPIN) Test

A material-specific objective intelligibility test, referred to as the Speech-in-Noise (SPIN) task, was done on a separate day after the MEG recordings. Due to the COVID-19 pandemic, only data from 32 subjects were obtained: 18 younger adults and 14 older adults. The task was run via from a graphical user interface in MATLAB. Subjects listened to 3– 5 s duration short sentence segments (with 4–7 key words) from the same audio book used for the MEG study, using different segments from those used in the MEG study (but processed identically). Participants were asked to repeat back the speech segment, in the case of quiet speech, and the selectively attended speech segmented, otherwise. At each noise condition there were six different speech segments; the first segment was used as a practice trial and was not included in the accuracy calculation. The accuracy per each condition and talker was calculated as the ratio (number of correctly repeated key words)/(total number of key words per condition and talker). The same conditions (quiet speech, 0 dB SNR, −6 dB SNR and babble speech; attend male and attend female) were used, in the same order presented as in the MEG study for that subject.

#### QuickSIN Test

The Quick Speech-in-Noise test (QuickSIN) (Killion et al. 2004), a standardized measure of a listener’s ability to understand speech in noise, was also employed. Due to the COVID-19 pandemic, only data from half of the subjects were obtained, and therefore these data were not further analyzed.

### Data analysis

#### Stimulus (Envelope) Reconstruction

Reconstruction of the speech envelope (backward/decoding model) from the neural response is a measure of cortical representation of the perceived speech. The low frequency envelopes from the attended (foreground; att) and unattended (background; unatt) talkers are denoted by *E_att_* (*t*) and *E_unatt_* (*t*) respectively. For each subject and each trial, the attended and unattended speech envelopes were reconstructed separately (but simultaneously) using a linear temporal decoder applied to the first 6 DSS components (*D*(*d*, *t*)) estimated by the Boosting algorithm (David et al. 2007; Ding and Simon 2013; Ding et al. 2014) as follows.

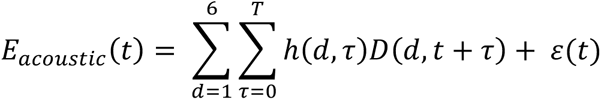

Where *ε*(*t*) is the contribution not explained by the model and ℎ(*d*, *τ*) is the decoder matrix value for component *d* at time lag *τ*. *T* is the integration window (500 ms unless specified otherwise). 10-fold cross-validation was used, resulting in 10 decoding filters per trial. These 10 decoders were averaged to produce the final decoding filter for each trial. This filter was then used to reconstruct the speech envelope, and the decoder accuracy is given by the linear correlation coefficient between reconstructed and the true speech envelope.

#### Integration Window Analysis

Performing reconstruction analysis with a fixed 500 ms integration window does not provide any access to temporal processing details within that window. Employing different time intervals allows incorporation of different information, and age-related differences in temporal processing should manifest as different trajectories for how envelope reconstruction accuracy builds up over time. Thus, the integration window size was also systematically varied from 10 ms to 610 ms with a step size of 50 ms. Generalized additive mixed models (GAMMs) were used to analyze the resulting time (integration window duration) series data.

#### Temporal Response Function (TRF)

Envelope reconstruction is a robust measure of how well a neural response tracks the stimulus, but any such backward model must necessarily integrate over information regarding neural response time and sensors (Haufe et al. 2014). In contrast, the TRF, which as a forward model relates how the neural responses were generated from the stimulus, allows interpretation of the stimulus-driven brain responses (Lalor et al. 2009; Ding and Simon 2012b), since it instead integrates over stimulus time, not response time.

TRF analysis, a linear method widely used to analyze the temporal processing of the auditory signal, predicts how the brain responds to acoustic features with respect to time. Additionally, a simultaneous two-talker TRF model uses the envelopes from both the foreground and background talkers, denoted by *E_att_* (*t*) and *E_unatt_* (*t*) respectively, with the model is formulated as:

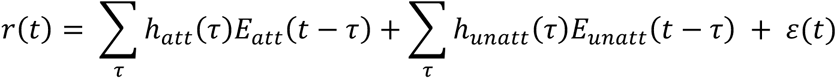

Where *r*(*t*) is the cortical response at a particular sensor, *τ* is the time lag relative to the speech envelope *E*(*t*), and *ε*(*t*) is the residual cortical response not explained by the linear model. ℎ*_att_* (*τ*) and ℎ*_unatt_* (*τ*), describe the filters that define the linear neural encoding from speech envelope to the neural response, and are known as the TRFs for the attended and unattended speech, respectively. The range of *τ* is chosen to range from 0 to 500 ms. The competing TRFs were estimated simultaneously using the boosting algorithm with 10-fold cross-validation, to minimize the mean square error between the predicted neural response and the true neural response (David et al. 2007; Zion Golumbic et al. 2013). For the babble condition, the summed three talker babble speech envelope was used as the background. The final TRF was evaluated as the mean TRF over the 10-fold cross-validation sets. TRFs were estimated for each condition and subject on the concatenated data giving one TRF per subject and condition (e.g., in the 0 dB case, all 6 such trials were concatenated before applying the boosting algorithm).

Larger amplitudes in the TRF indicate that the neural populations with the corresponding latencies follow the speech envelope better when synchronously responding to the stimulus. The TRF has three prominent peaks, with latencies at ∼50 ms (positive peak), ∼100 ms (negative peak) and 200 ms (positive peak), named as the M50_TRF_, M100_TRF_ and M200_TRF_ respectively. Each peak corresponds to a different stage in the auditory signal processing chain. Latency and polarity of these peaks under different conditions can be compared similarly to the P1, N1 and P2 CAEP peaks. For example, an age- or task-related increase in M50_TRF_ amplitude represents a stronger response at that latency under identical stimulus conditions. For each subject and condition, peak latencies were extracted within a specific time range; for M50_TRF_, M100_TRF_ and M200_TRF_ the windows were 30–110 ms, 80–200 ms and 140–300 ms respectively. These peak amplitudes and latencies were further analyzed to evaluate the effects of aging, task difficulty, and selective attention.

### Statistical analysis

All statistical analysis was performed in R version 4.0 (R Core Team 2020). Linear mixed effect models (LMM) were used to systematically evaluate the relationships between the dependent (behavioral scores, neural measures) and independent variables (age, noise condition, selective attention). For the LMM analysis, the *lme4* (version 1.1-30) (Bates et al. 2015) and *lmerTest* (version 3.1-30) (Kuznetsova et al. 2017) packages in R were used. The best fit model from the initial full model was found by the *buildmer* package (version 2.4) (Voeten 2020) using the default settings, where the *buildmer* function first determines the order of parameters based on the likelihood-ratio test (LRT) and then uses a backwards elimination stepwise procedure to identify the model of best fit for random and fixed effects. The assumptions of mixed effect modelling, linearity, homogeneity of variance and normality of residuals, were checked per each best fit model based on the residual plots. Reported *β* values represent changes in the dependent measure when comparing one level of an independent variable versus its reference level. p-values were calculated using Satterthwaite approximation for degrees of freedom (Satterthwaite 1941; Luke 2017). In order to interpret significant fixed effect interaction terms, variables were releveled to obtain model estimates for individual factor levels (indicated by ‘with [*new reference level*] reference level’). The summary tables for each model used in result section are reported in the *Appendix*.

The subjective intelligibility ratings and objective intelligibility scores (SPIN scores) were analyzed separately, each using a LMM with *age* as a between-subject factor (categorical variable; 2 levels: younger [reference level], older) and *noise condition* as a within-subject factor (categorical variable; 4 levels: quiet [reference level], 0 dB, −6 dB and babble). To account for repeated measures, we used *subject* as a clustering variable so that the intercept and effects of *noise condition* could vary across subjects (random intercept for *subject* and random slopes for *noise condition* by *subject* respectively). The full model for each dependent variable was defined as *intelligibility ∼ age × noise condition +* (*noise condition |subject*). To evaluate the relationship between the two measures, data were analyzed using a separate LMM with *SPIN score* as the dependent variable, intelligibility rating as the independent variable and *subject* as a random intercept: SPIN score *∼ intelligibility rating +* (*1|subject*).

LMMs were used to systematically evaluate the relationships among the computed neural features (reconstruction accuracy; M50_TRF_, M100_TRF_, M200_TRF_ for both amplitude and latency) and *age*, *noise condition*, and selective *attention* (categorical variable; 2 levels: attended [reference level], unattended). Two models were generated for each neural feature, 1) to examine the effects of aging and noise condition on the neural features measured for the attended talker, 2) to examine the effects of aging, noise condition and attention on the neural features measured for the attended talker and unattended talker. Only data from two competing talkers noise conditions were used for the latter model. The full models for 1) and 2) were defined as*: neural feature ∼ age × noise condition +* (*1 + noise condition |subject*) and *neural feature ∼ age × noise condition × attention +* (*1 + noise condition × attention |subject*), respectively.

To model nonlinear changes in the integration window analysis, Generalized Additive Mixed Models (GAMMs) (Wood 2006) in R (packages *mgcv (version 1.8-40)*, *itsadug (version 2.4.1)*) were used to analyze the reconstruction accuracies over integration window. Compared to Generalized Linear Mixed Effect Models, GAMMs have several advantages for modeling time series data, especially in electrophysiology (DeCat et al. 2014; Tremblay and Newman 2015) and pupillometry (van Rij et al. 2019). In particular, it 1) can model both linear and non-linear patterns in the data using *parametric* and *smooth terms* and 2) can, critically, include various types of autoregressive (AR) correlation structures that deal with autocorrelational structure in the errors. Compared to conventional non-linear mixed effect modelling where non-linear trends are fitted by polynomials of the predictor, GAMMs fit with p-spline based “smooth terms” with a specified number of basis functions (knots) that specify how “wiggly” the model can be. For each fixed effect term (age, noise condition and attention) and corresponding smooth term, model testing was compared between a test model and baseline model using Chi-Square (*compareML* in the *itsadug* package) to determine the significance of predictors (Sóskuthy 2017; van Rij et al. 2019). The parametric term indicates the overall difference in height between two curves, whereas smooth terms indicate the difference in shape (i.e., “wiggliness”) between two curves. The models included random smooths for each subject to capture the individual trends in reconstruction accuracies over integration window. For the time series reconstruction accuracies here, since the current time point was observed to be correlated with the next time point, we employed an autoregression model 1 (AR1) structure. The assumptions of mixed effect modelling and model diagnostics (over-smooth or under-smooth) were performed by residual plots and the *gam.check()*function.

### Code and data availability

Preprocessed MEG data, stimulus material, behavioral data, and analysis codes are available at https://doi.org/10.13016/4goq-vs0z

## RESULTS

### Behavioral Data

#### Flanker test

The effect of age on the flanker scores was analyzed with a two-sample t-test. Results showed significantly better scores in younger adults than for older adults (*t*_32_ = 6.0956, *p* < 0.001), suggesting that older adults’ performance in inhibition task may decline with aging.

#### SPIN test

The effects of age and noise condition on the objective intelligibility measures (SPIN scores) were analyzed using LMM. Figure 2(a) plots the SPIN scores for both groups at all noise conditions. The best fit model included main effects of noise condition on speech SPIN scores and random intercept by subject (Table A1). There was no effect of age on the SPIN scores. SPIN scores significantly dropped from quiet speech to every other condition (Quiet to 0 dB to −6 dB to Babble) with the highest drop from 0 dB to −6 dB (with Quiet reference level: *noise condition*(0 dB): *β* = −12.88, *SE* = 1.57, *p* < 0.001; with 0 dB reference level: *noise condition*(−6 dB): *β* = - 40.95, *SE* = 1.57, *p* < 0.001; with −6 dB reference level: *noise condition*(Babble): *β* = −13.52, *SE* = 1.92, *p* < 0.001). The lack of significant age effects will be addressed in discussion.

**Figure 2.**
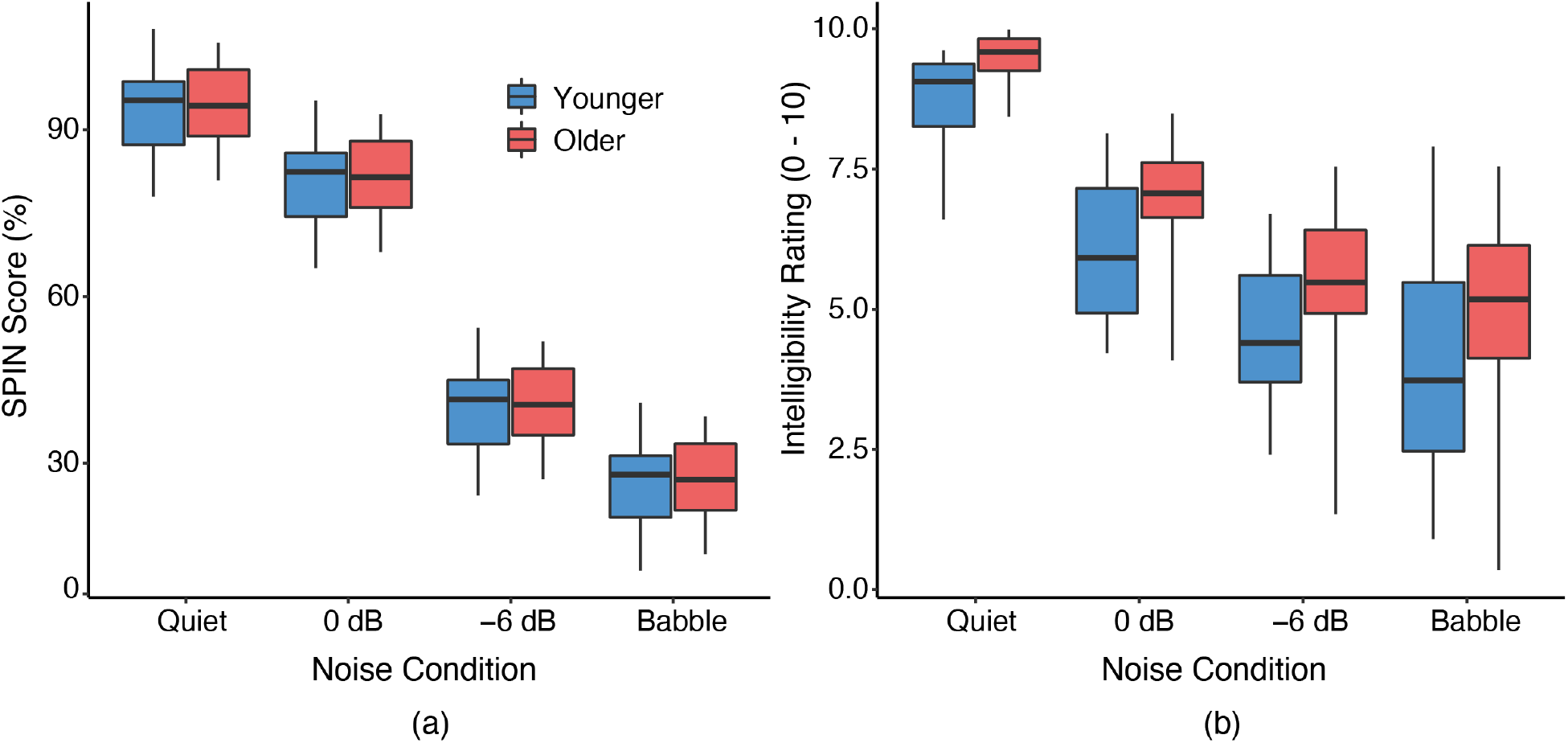
Model predicted behavioral test results for (a) speech SPIN scores (0–100%) and (b) intelligibility ratings (0–10). Both scores drop significantly with the noise condition. A significant effect of age was seen only for the intelligibility rating whereas no age effect was found on the SPIN scores

#### Intelligibility ratings

Parallel LMM analysis of the subjective intelligibility ratings Figure 2(b) revealed fixed effects of both *age* and *noise condition* along with random slopes for *noise condition* by *subject* (Table A2). Older adults rated the intelligibility slightly higher compared to younger adults (*β* = 0.76, *SE* = 0.36, *t* = 2.11, *p* = 0.04). As in the case of SPIN scores, intelligibility ratings dropped significantly from quiet speech to 0 dB SNR to −6 dB to Babble noise condition in both groups (with Quiet reference level: *noise condition*(0 dB): *β* = −2.6, *SE* = 0.27, *p* < 0.001, with 0 dB reference level: *noise condition*(−6 dB): *β* = −1.51, *SE* = 0.27, p < 0.001, with −6 dB reference level: *noise condition*(Babble): *β* = −0.53, *SE* = 0.34, *p* = 0.13).

To examine the consistency of the two measures, a separate LMM model was constructed to predict SPIN scores from intelligibility ratings. The best fit model revealed that the subjective intelligibility ratings were positively related to the objective intelligibility scores (*β* = 8.2, *SE* = 0.58, *p* < 0.001) and revealed no effects of age.

### Stimulus (Envelope) Reconstruction Analysis

As a simpler precursor to the full TRF analysis, we employed reconstruction analysis to investigate how the cortical representation of the speech envelope is affected by aging at a coarser level. First, we investigated the effects of age and noise condition on the attended talker envelope tracking. As summarized in Table A3 the best fit model revealed main effects of *age*, *noise condition* and *age* × *noise condition* interactions with random intercepts by subject. For both groups and for all noise conditions, the reconstruction accuracies fitted by the model for the attended talker are shown in Figure 3a. The main effects of *age* revealed that aging is associated with higher reconstruction accuracies (*β* = 0.04, *SE* = 0.01, *p* < 0.001) in all noise conditions.

**Figure 3.**
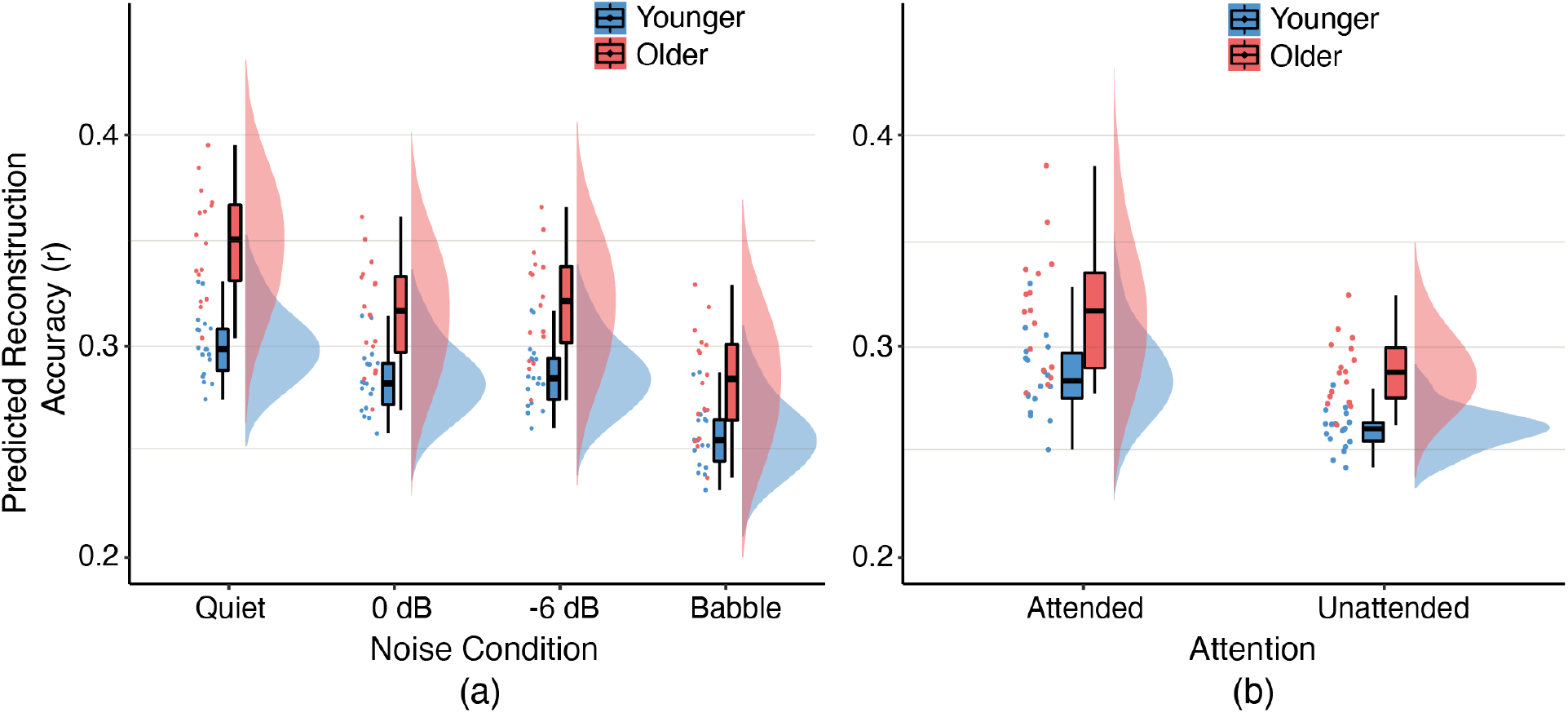
Model-predicted values for reconstruction accuracy for (a) the attended speech for both younger and older adults and for all noise conditions, and (b) the attended vs unattended speech envelope reconstruction accuracies for competing talker conditions (there is no separation by noise condition since no significant dependence on noise condition was found). Both attended and unattended speech envelope reconstruction illustrates that the speech reconstruction was significantly higher in older adults. When a single or multiple competing talkers are added, attended talker reconstruction accuracies significantly decrease in both groups. In both groups attended talker envelope reconstruction was higher compared to unattended.

The significant interaction *age* × *noise condition* term revealed that the aging adversely affects speech reconstruction from quiet to noisy speech (*age*(Older)×*noise condition*(0 dB): *β* = −0.018, *SE* = 0.01, *p* = 0.01, *age*(Older)×*noise condition*(−6 dB): *β* = −0.015, *SE* = 0.01, *p* = 0.03, *age*(Older)×*noise condition*(Babble): *β* = −0.023, *SE* = 0.01, *p* = 0.01). However, this effect was not significant from noisy speech to babble speech (with 0 dB reference level: *age*(Older)×*noise condition*(Babble): *β* = −0.005, *SE* = 0.01, *p* = 0.58). As can be seen from Figure 3a, the main effect of noise condition revealed that the attended talker envelope reconstruction accuracies significantly reduce from quiet to noisy conditions in both groups (*noise condition*(0 dB): *β* = - 0.02, *SE* = 0.004, p < 0.001, *noise condition*(−6 dB): *β* = −0.02, *SE* = 0.004, *p* < 0.001, *noise condition*(Babble): *β* = −0.05, *SE* = 0.005*, p* < 0.001). No significant difference was observed between the 0 dB and −6 dB noise condition (with 0 dB reference level: *noise condition*(−6 dB): *β* = 0.003, *SE* = 0.004, *p* = 0.39). However, reconstruction accuracies significantly dropped from noisy speech to babble speech (with 0 dB reference level: *noise condition*(Babble): *β* = −0.03, *SE* = 0.005, *p* < 0.001, with −6 dB reference level: *noise condition*(Babble): *β* = −0.03, *SE* = 0.005, *p* < 0.001).

Secondly, to investigate the combined effects of selective attention, age, and noise condition, a separate analysis was performed on only the competing talker speech data (0 dB and −6 dB) by including both attended and unattended speech envelope reconstruction accuracies. As shown in Figure 3b, LMM analysis revealed main effects of *age* and *selective attention* on reconstruction accuracy, with both random intercepts and random slopes for *selective attention*, by *subject* (Table A4). Results revealed that the cortical representation of the speech envelope as measured by reconstruction accuracy is enlarged/overrepresented in older adults for both attended and unattended talkers (*β* = 0.03, *SE* = 0.01, *p* < 0.001). Furthermore, in both groups the attended talker was better represented than the unattended talker (*β* = −0.03, *SE* = 0.004, *p* < 0.001). For attended talker vs. unattended talker, or for either of the age groups, no significant difference was observed between the 0 dB and −6 dB noise conditions.

### Integration Window Analysis

Integration window analysis was done using GAMMs including both quiet and two talker mixed speech for both attended and unattended speech envelope reconstruction accuracies. The initial model included a smooth term over the integration window, characterizing the nonlinearity of these functions. Model comparisons determined that separate smooths for *age* × *noise condition* × *attention* significantly improve the model fit (*χ*^2^(23) = 212.1, *p* < 0.001). Moreover, adding the fixed effect term, characterizing the height of these functions, *age* × *noise condition* × *attention*, and random smooth per subjects, also improved the model fit (*χ*^2^(32) = 1531.39, *p* < 0.001, *χ*^2^(34) = 1886.42, *p* < 0.001 respectively). Finally, as the residual plots showed a high autocorrelation in the residual analysis, an autoregression (AR1) model was included. The statistical information on this model (parametric terms and smooth terms) is summarized in the Appendix (Table A10). Figure 4(a) shows the resulting smooth plots for the two groups. The results show that envelope reconstruction accuracy initially rapidly increases as the time window duration increases, and then stabilizes to a slower rate as longer latencies are included.

**Figure 4.**
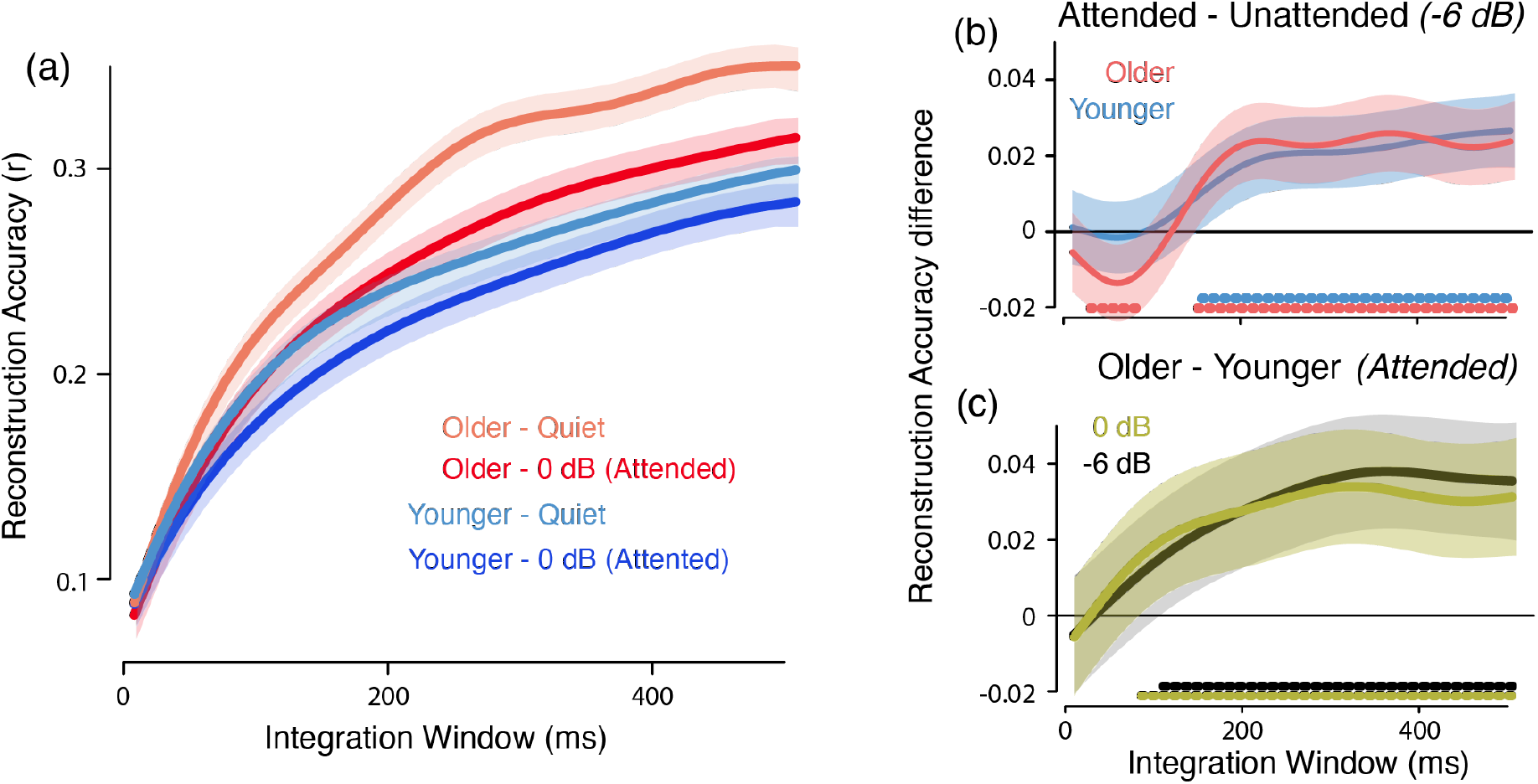
Integration Window Analysis using GAMMs. (a) Changing reconstruction accuracy with integration window (only a subset of curves is shown, for visual clarity). (b) Reconstruction accuracy difference between attended vs unattended talker for −6 dB noise condition. (c) Reconstruction accuracy difference between older vs younger for attended talker. The color-coded horizontal lines above the horizontal axis in (b, c) mark where the difference is statistically significant. Shaded area represents the 95% confidence interval (CI). In both groups reconstruction accuracy initially increases rapidly with increasing integration window, slowing down after ∼300 ms. The attended talker is better represented than the unattended after ∼170 ms. The overrepresentation of attended talker envelope starts at early processing stages (<100 ms).

To investigate how the integration window affects selective attention effects, the difference between attended and unattended talker responses were analyzed. Figure 4(b) shows the dynamical differences between attended and unattended talker reconstruction accuracy curves for both younger and older adults and for the 0 dB noise condition. The color-coded horizontal lines at graph bottom indicate where the differences are significant. In both age groups, attended talker reconstruction accuracies were significantly higher compared to unattended talker after the middle processing stage (∼150 ms). Interestingly, in older adults, for the −6 dB noise condition, the unattended talker representation is enhanced compared to the attended talker during the early processing stages (∼50 ms). Difference analysis emphasizing age-group differences revealed that older adults’ overrepresentation of the speech envelope starts as early as ∼85 ms for the attended (Figure 4(c)) and 55 ms for the unattended talker when averaged across noise conditions. As can be seen from Figure 4(c), the difference monotonically increases until ∼300 ms and then levels off, suggesting that older adults rate of increase in reconstruction accuracy with integration window is higher compared to younger adults until later processing stages.

### Temporal Response Function Analysis

Unlike stimulus reconstruction analysis, which integrates over latencies, TRF analysis allows direct analysis of neural processing associated with any latency. For each TRF component (M50_TRF_, M100_TRF_ and M200_TRF_) we separately analyzed individuals’ peak amplitude and latency, comparing between the two age groups, and for both attended and unattended talker and noise conditions. Figure 5(a) visualizes how the TRF peak amplitudes and latencies varied for the attended talker across two age groups and noise conditions. Figure 6(a) visualizes how the TRF peak amplitudes and latencies varied between attended vs unattended talker.

**Figure 5.**
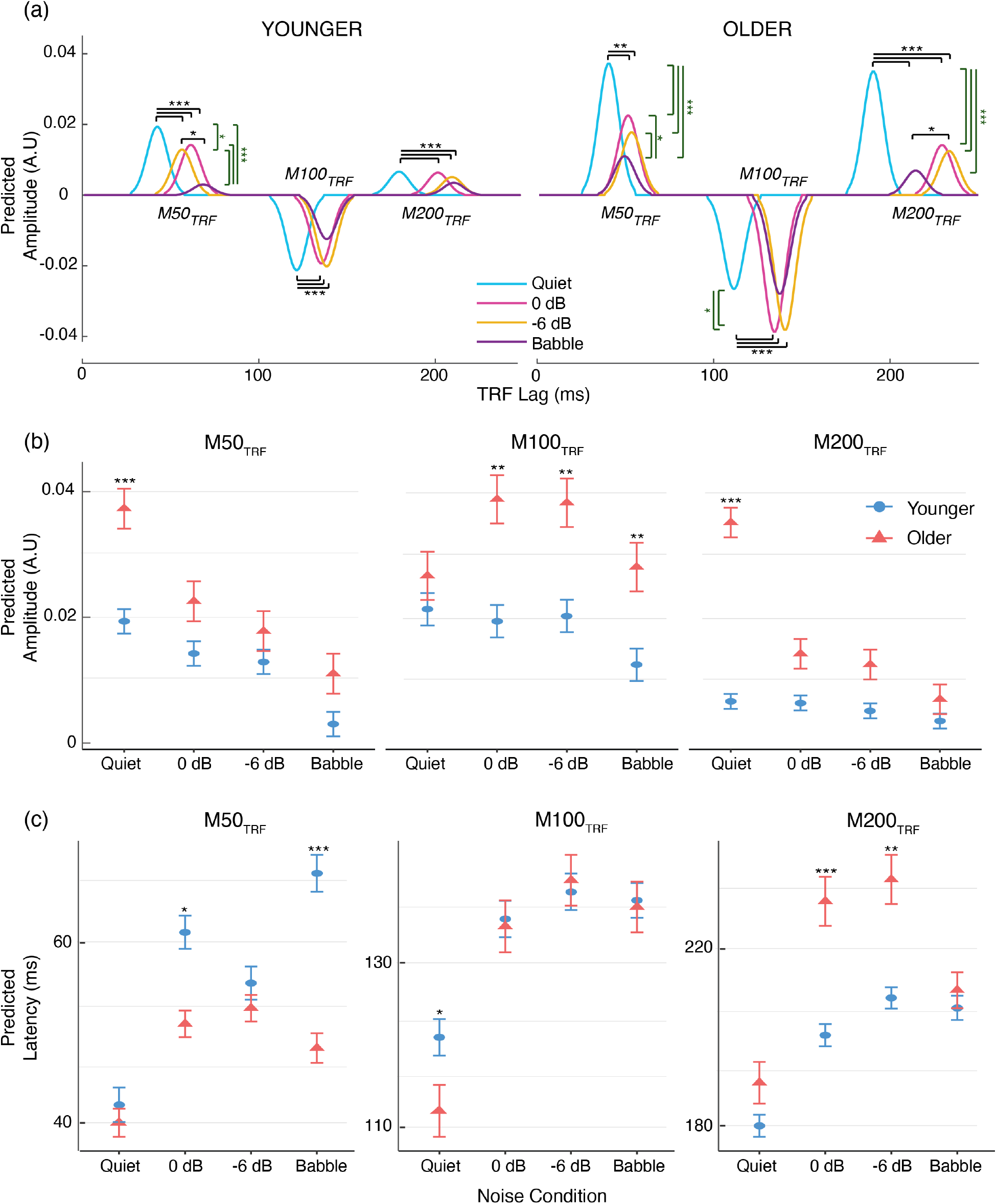
Model-predicted attended talker TRF peak amplitudes and latencies. (a) TRFs showing overall amplitudes and latency for both groups and all noise conditions (for visualization simplicity, all peaks are represented with the same Gaussian shape, standard deviation 7 ms, centered at the group mean peak latency, and with amplitude given by the group mean peak amplitude). (b) The TRF amplitudes (±SE) as a function of noise condition (for the M100_TRF_, as a negative polarity peak, the unsigned magnitude is shown). Generally, older adults (red) exhibit stronger TRF peak amplitudes compared to younger adults (blue). When a competing talker is added to the stimulus, the attended M50_TRF_ amplitudes decrease in both groups. From the quiet to competing speech conditions, the M100_TRF_ increases and M200_TRF_ decreases but only in older adults. (c) The TRF latencies (±SE) as a function of noise condition. Compared to younger adults, in older adults the M50_TRF_ is earlier and the M200_TRF_ is later. With task difficulty, peaks are typically delayed in both groups with some exceptions in the babble condition. *p<0.05, **p<0.01, ***p<0.001

**Figure 6.**
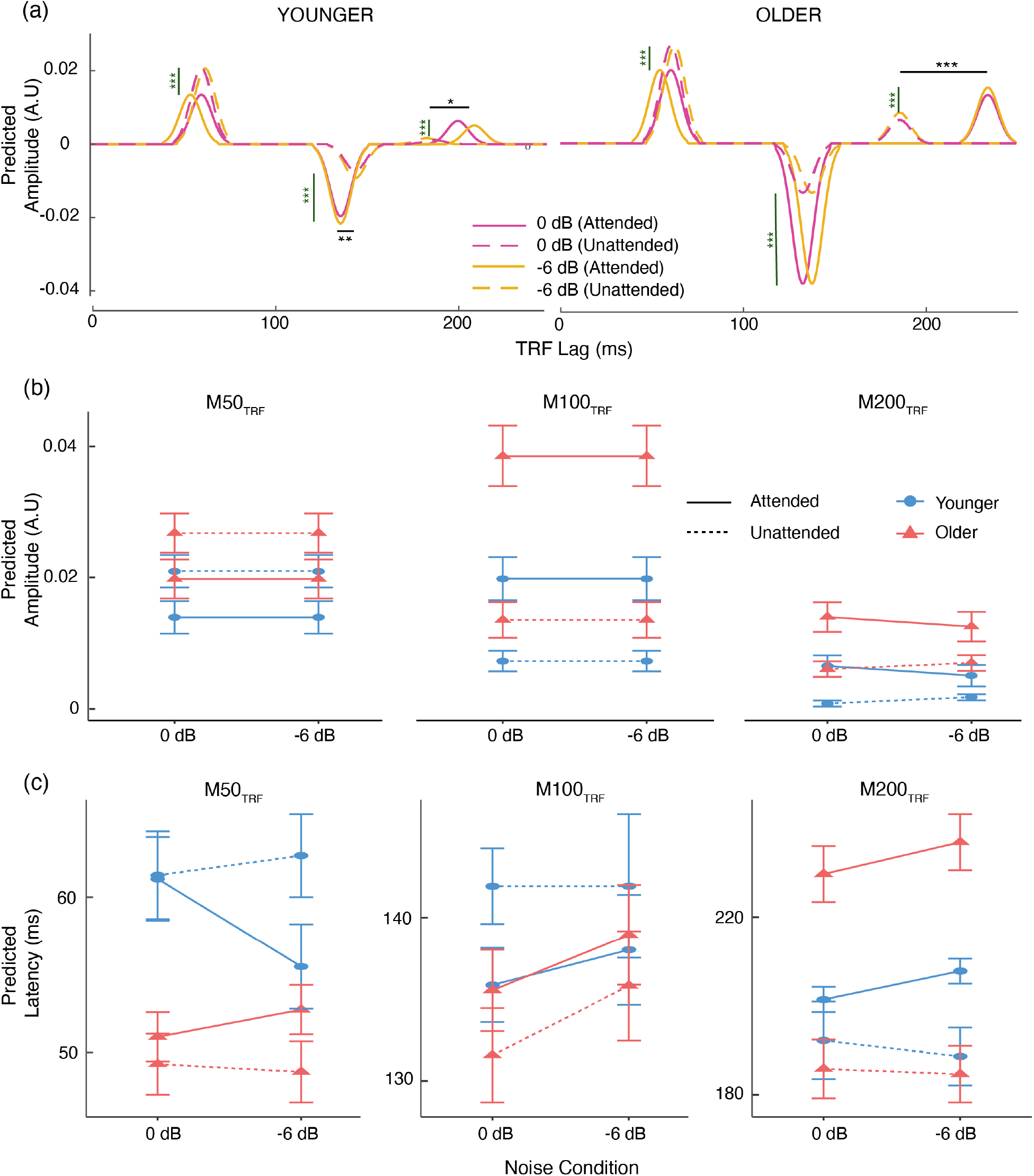
Model-predicted values for attended vs unattended talker TRF peak amplitude and latency. (a) TRF peak amplitudes and latency for both groups and two talker speech conditions for both attended and unattended talker (solid line = Attended, dashed line = Unattended). For visualization simplification, peaks are represented with a common Gaussian shape as in Figure 5. (b) TRF amplitudes (±SE) as a function of noise condition (for the M100_TRF_, as a negative polarity peak, the unsigned magnitude is shown). Generally older adults exhibit stronger TRF amplitudes for both attended and unattended peaks. Attended M50_TRF_ is significantly smaller compared to unattended amplitudes. In contrast, attended M100_TRF_ and M200_TRF_ are enhanced compared to unattended peak amplitude. Note that the model analysis for the M50_TRF_ and M100_TRF_ amplitude did not find significant differences between noise conditions, so the model mean and SE do not change there. (c) TRF latencies (±SE) as a function of noise condition. The attended M200_TRF_ peak is significantly delayed compared to the unattended M200_TRF_ peak and this difference is bigger in older adults. *p<0.05, **p<0.01, ***p<0.001

#### TRF peak amplitudes

LMMs were fitted to the M50_TRF_, M100_TRF_ and M200_TRF_ amplitudes separately to analyze the effect of age and noise condition on the attended talker TRF peak amplitudes for each neural processing stage. The best fit model indicated main effects of *age, noise condition* and *age × noise condition* interaction along with random intercept by subject for all three peaks M50_TRF,_ M100_TRF_, and M200_TRF_ (Table A5).

TRF peak amplitudes for the M50_TRF_, M100_TRF_ and M200_TRF_ are plotted in Figure 5(a, b). Overall, in the comparison between younger vs older, older adults showed exaggerated peak amplitudes in all 3 processing stages (M50_TRF_: *age*(Older): *β* = −0.01, *SE* = 0.004, *p* = 0.03, M100_TRF_: *age*(Older): *β* = −0.02, *SE* = 0.005, *p* = 0.008 and M200_TRF_: *age*(Older): *β* = −0.018, *SE* = 0.003, *p* = 0.001). For both the M50_TRF_ and M200_TRF_, peak amplitudes were stronger in all noise conditions and that was significant only for the quiet speech condition (M50_TRF_: *age*(Older): *β* = 0.02, *SE* = 0.005, *p* < 0.001, M200_TRF_: *age*(Older): *β* = 0.03, *SE* = 0.01, *p* < 0.001). In contradistinction, except for the quiet speech, the M100_TRF_ was stronger in all the noisy conditions (*age*(Older): *β =* 0.01, *SE* = 0.01, *p* = 0.036, with 0 dB reference level: *age*(Older): *β* = 0.02, *SE* = 0.01, *p* = 0.001, with −6 dB reference level: *ag*e(Older): *β* = 0.02, *SE* = 0.01, *p* = 0.002, with Babble reference level: *age*(Older): *β* = 0.02, *SE* = 0.01, *p* = 0.008).

The effects of noise condition revealed that the M50_TRF_ response decreases from quiet to every other noise condition in both groups (*noise condition*(0 dB): *β* = −0.01, *SE* = 0.003, *p* = 0.01, *noise condition*(−6 dB): *β* = −0.01, *SE* = 0.003, *p* = 0.04, *noise condition*(Babble): *β* = −0.02, *SE* = 0.003, *p* < 0.001, with Older reference level: *noise condition*(0 dB): *β* = −0.01, *SE* = 0.003, *p* < 0.001, *noise condition*(−6 dB): *β* = −0.2, *SE* = 0.003, *p* < 0.001, *noise condition*(Babble): *β* = −0.03, *SE* = 0.003, *p* < 0.001). A significant *age* × *noise condition* interaction indicated that aging contributes more to the M50_TRF_ reduction as a function of noise condition (*age*(Older) ×*noise condition*(0 dB): *β* = −0.01, *SE* = 0.004, *p* = 0.03). In contrast to the M50_TRF_, the M100_TRF_ and M200_TRF_ did not significantly vary across the quiet and two talker noisy conditions in younger adults. However, in older adults, from quiet to two-talker noise conditions the M100_TRF_ significantly increased (with Older reference level: *noise condition*(0 dB): *β* = 0.01, *SE* = 0.003, *p* < 0.001), while the M200_TRF_ decreased (with Older reference level: *noise condition*(0 dB): *β* = −0.02, *SE* = 0.003, *p* < 0.001). Interestingly, in both groups the M100_TRF_ peak amplitudes significantly dropped from −6 dB to the babble condition (with −6 dB reference level: *noise condition*(Babble): *β* = −0.01, *SE* = 0.003, *p* = 0.02, with Older, −6 dB reference level: *noise condition*(Babble): *β* = −0.01, *SE* = 0.003, *p* = 0.003).

To investigate the combined effects of aging, selective attention and noise condition on the TRF amplitude responses, separate LMM models were constructed (Table A6). Figure 6(b) displays the TRF amplitude variation for the attended and unattended talker, for both age groups and for competing talker conditions (0 dB and −6 dB). LMM applied to the M50_TRF_ showed a main effect of *attention*, revealing that the unattended M50_TRF_ amplitude is bigger compared to the attended M50_TRF_ amplitude in both groups (*attention*(Unattended): *β* = 0.01, *SE* = 0.001, *p* < 0.001). In contrast, the M100_TRF_ peak amplitudes showed main effects of *age*, *attention* and an *age* × *attention* interaction effect. Compared to younger adults, both the attended and unattended M100_TRF_ peak amplitudes were exaggerated in older adults, however this effect was statistically significant only for the attended talker peak amplitudes (*age*(Older): *β* = 0.02, *SE* = 0.01, *p* < 0.001, with Unattended reference level: *attention*(Unattended): *age*(Older): *β* = 0.01, *SE* = 0.01, *p* = 0.19). In both age groups unattended peak amplitudes were reduced compared to attended peak amplitudes (*attention*(Unattended): *β* = −0.01, *SE* = 0.003, *p* < 0.001, with Older reference level: *attention*(Unattended): *β* = −0.02, *SE* = 0.003, *p* < 0.001), and the interaction effect revealed that this reduction in peak heights is amplified by aging (*β* = −0.01, *SE* = 0.004, *p* < 0.001). Interestingly, the M200_TRF_ showed main effects of *age* and *attention*, and an *attention* × *noise condition* interaction. M200_TRF_ amplitudes in both attended and unattended TRFs were stronger in older adults (*age*(Older): *β* = 0.01, *SE* = 0.001, *p* < 0.001). The selective attention effect revealed that in both groups, the attended talker M200_TRF_ peak amplitude is stronger compared to the unattended (*attention*(Unattended): *β* = −0.01, *SE* = 0.002, *p* < 0.001).

#### TRF peak latencies

Similar to TRF peak amplitude analysis, TRF peak latency analysis was performed on the attended talker TRFs as the first step. The best fit model revealed effects of *age*, *noise condition* and *age* × *noise condition* along with random intercepts by subject for M50_TRF_, M100_TRF_ and M200_TRF_ (Table A7). Model predicted latencies are plotted in Figure 5(c). Averaged latencies over noise conditions revealed that compared to younger adults, the older adults’ early peak, M50_TRF_, is significantly earlier (*age*(Older): *β* = 8.4, *SE* = 3.1, *p* = 0.01) and that there is no significant latency difference for the middle peak, M100_TRF_ (*age*(Older): *β* = 2.46, *SE* = 4.15, *p* = 0.55) whereas the late peak M200_TRF_ is significantly delayed (*age*(Older): *β* = −17.2, *SE* = 6.37, *p* = 0.01). These results suggest that the three distinct processing stages are each differently affected by aging.

All three peaks were significantly delayed for the noisy conditions, relative to quiet, for both younger (M50_TRF_ : *noise condition*(0 dB): *β* = 18.48, *SE* = 3.02, *p* < 0.001, M100_TRF_ : *noise condition*(0 dB): *β* = 14.59, *SE* = 2.75, *p* < 0.001, M200_TRF_ : *noise condition*(0 dB): *β* = 21.80, *SE* = 4.64, *p* < 0.001) and older adults (with Older reference level: M50_TRF_ : *noise condition*(0 dB): *β* = 10.83, *SE* = 3.09, *p* < 0.001, M100_TRF_ : *noise condition*(0 dB): *β* = 22.13, *SE* = 2.92, *p* < 0.001, M200_TRF_ : *noise condition*(0 dB): *β* = 38.80, *SE* = 4.86, *p* < 0.001) highlighting that the peak responses are delayed with the stimulus noise condition. With respect to quiet, babble speech latencies in all three processing were delayed in both younger (*noise condition*(Babble): M50_TRF_ : *β* = 25.62, *SE* = 3.12, *p* < 0.001*, M100_TRF_ : β* = 16.99, *SE* = 2.71, *p* < 0.001, M200_TRF_ : *β* = 30.89, *SE* = 5.17*, p* < 0.001) and older adults (with Older reference level: *noise condition*(Babble): M50_TRF_ : *β* = 9.24, *SE* = 3.23, *p* = 0.006, M100_TRF_ : *β* = 25.63, *SE* = 2.87, *p* < 0.001, M200_TRF_ : *β* = 24.59, *SE* = 4.98, *p* < 0.001). The *age* × *noise condition* interaction manifests as aging contributing more to the peak delay from quiet to 0 dB for both M100_TRF_ (*age*(Older) *× noise condition*(0 dB)*: β* = 8.05, *SE* = 3.98, *p* = 0.04) and M200_TRF_ (*age*(Older) *× noise condition*(0 dB): *β* = 17.08, *SE* = 6.68, *p* = 0.01).

The effect of selective attention on TRF peak latencies was analyzed using LMMs (Table A8) and results are displayed in Figure 6(c). Comparing mean latencies across noise conditions revealed that compared to younger adults, unattended peaks are earlier in older adults for both the M50_TRF_ and M100_TRF_ (with Unattended reference level: *age*(Older): M50_TRF_: *β* = −12.96, *SE* = 3.96, *p* < 0.001, M100_TRF_: *β* = −13.98, *SE* = 4.23, *p* = 0.003). This effect was, however, not significant for the M200_TRF_ (with Unattended reference level: *age*(Older): *β* = −0.70, *SE* = 8.65, *p* = 0.94). Differences due to selective attention on the neural response latencies were analyzed with the same LMM models. Results indicated that there is no significant latency difference between attended and unattended peaks for the early peak M50_TRF_ in both groups (*attention*(Unattended): *β* = 0.50, *SE* = 3.11, *p* = 0.87, with Older reference level: *attention*(Unattended): *β* = 2.88, *SE* =2.83, *p* = 0.32), whereas for the middle peak, M100_TRF_, the attended peak was earlier compared to unattended only in younger adults (*attention*(Unattended): *β* = 8.03, *SE* = 3.21, *p* = 0.01). For the late peak M200_TRF_, both age groups showed a delayed attended peak compared to the unattended peak (*attention*(Unattended): *β* = −12.02, *SE* = 6.28, *p* = 0.05), and this effect was stronger in older adults (*age*(Older)*×attention*(Unattended): *β* = −28.54, *SE* = 8.26, *p* = 0.001).

#### Amplitude vs Latency analysis

Potential associations between TRF peak amplitudes and latencies for the attended talker were analyzed for M50_TRF_, M100_TRF_ and M200_TRF_ separately. The analysis was conducted on 0 dB and −6 dB noise conditions (quiet and babble conditions were excluded, as they represent two extreme acoustic environments resulting in heavily restricted dynamic range in both amplitude and latency). Peak amplitudes were predicted by age and peak latencies. As can be seen from Figure 7, older adults’ M200_TRF_ peak amplitudes exhibited a significant negative relationship with the latency (with Older reference level: *Latency*: 0 dB condition: *β* = −0.0001, *SE* = 0.0001, *p* = 0.02, −6 dB condition: *β* = −0.0002, *SE* = 0.0001, *p* = 0.01), i.e., delayed peaks showed smaller peak amplitudes, but this was not seen for the earlier peaks. Conversely, in younger adults peak amplitudes were not related to the latencies (*Latency*: 0 dB condition: *β* = 0.0002, *SE* = 0.0001, *p* = 0.1, −6 dB condition: *β* = 0.0003, *SE* = 0.0001, *p* = 0.1) (Table A9).

**Figure 7.**
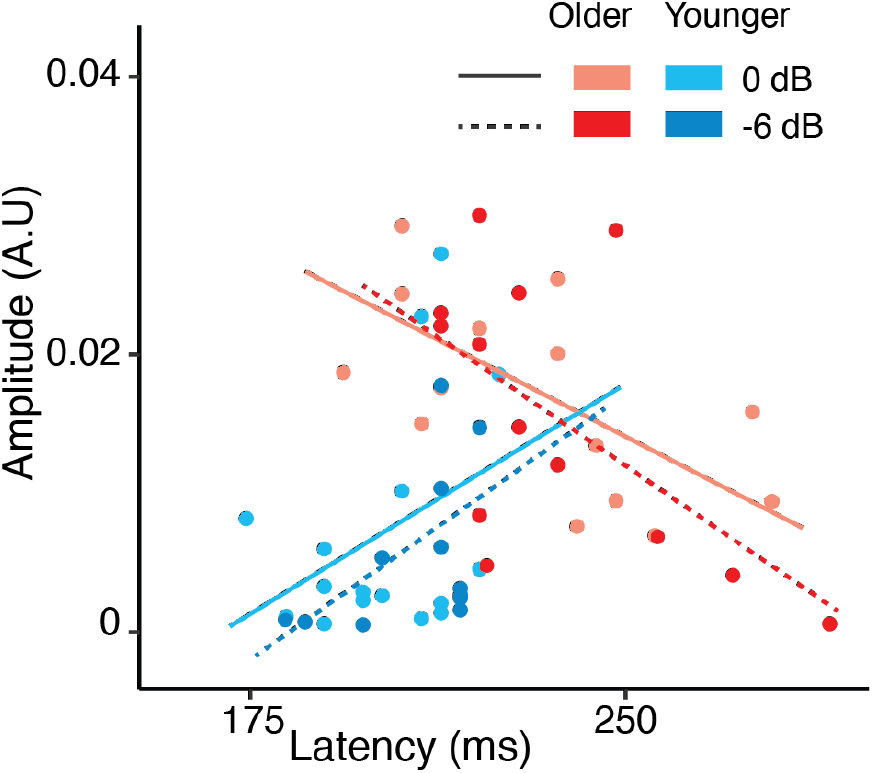
M200_TRF_ peak amplitude vs latency for 0 dB and −6 dB conditions. Older adults’ amplitudes were significantly negatively associated with their latencies; younger adults showed no significant association.

### Relationships among neural features and behavioral responses

LMMs were used to evaluate the relationship between the neural measures (reconstruction accuracies, TRF peak amplitudes and latencies) and the behavioral measures. Firstly, we analyzed how attended talker neural features are affected by speech intelligibility score and age but without specific regard to stimulus noise condition (noise condition and behavioral intelligibility measures are too correlated to include both measures, See Figure 2(a)). Results revealed that the reconstruction accuracy increases with better speech intelligibility in both groups (*SPIN score*: *β* = 0.0005, *SE* = 0.001, *p* < 0.001) (Figure 8(a)). Similarly, TRF peak amplitude analysis revealed that stronger M50_TRF_ amplitudes are associated with better speech intelligibility score (*SPIN score*: *β* = 0.0002, *SE* = 0.001, *p* < 0.001) (Figure 8(b)), and this effect was stronger in older adults (*age*(Older)*×SPIN score*: *β* = 0.0002, *SE* = 0.001, *p* = 0.01). However, no significant trends were found for M100_TRF_ amplitude. Interestingly, for the late peak M200_TRF,_ older adults showed smaller peak amplitudes with poorer speech intelligibility score (with Older reference level: *SPIN score*: *β* = 0.0003, *SE* = 0.001, *p* < 0.001), whereas no significant trend was found for younger adults (*SPIN score*: *β* = 0.0001, *SE* = 0.001, *p* = 0.022) (Figure 8(c)). Analysis of peak latencies revealed that, in both groups, peak latencies at all three stages are negatively related to the speech intelligibility score (*SPIN score*: M50_TRF_: *β* = −0.17, *SE* = 0.04, *p* < 0.001, M100_TRF_: *β* = −0.22, *SE* = 0.03, *p* < 0.001, M200_TRF_: *β* = −0.32, *SE* = 0.06, *p* < 0.001). Additionally, similar trends were observed when subjective intelligibility rating was used instead the SPIN scores.

**Figure 8.**
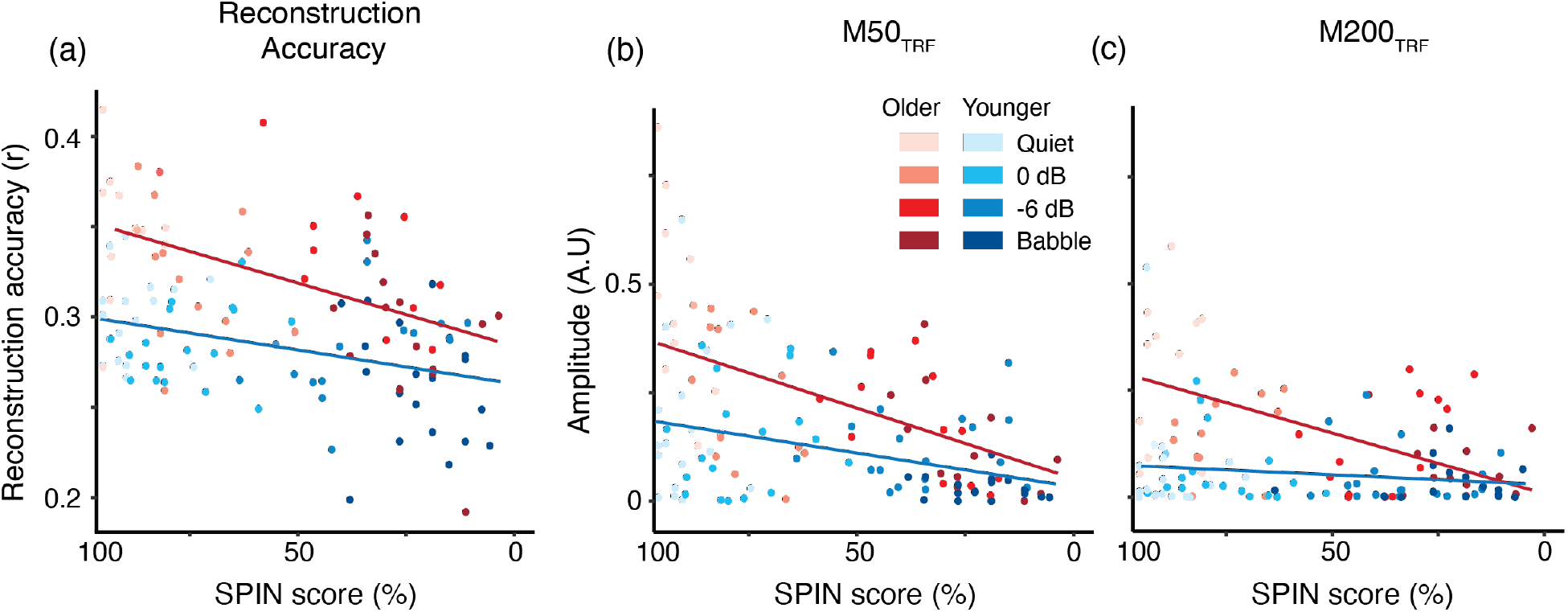
Neural measure vs SPIN scores including all noise conditions. (a) Reconstruction accuracy vs SPIN score. (b) M50_TRF_ peak amplitude vs SPIN score. Better reconstruction accuracies or M50_TRF_ peak amplitudes were related to better speech intelligibility scores in both groups. (c). M200_TRF_ peak amplitude vs SPIN score. Only in older adults, stronger M200_TRF_ peak amplitude was associated with better speech intelligibility score.

However, when noise condition was added into the model, for any one noise condition, no consistent trends were found between behavioral scores and neural measures.

## DISCUSSION

This study examined the effects of aging on neural measures of cortical continuous speech processing under difficult listening conditions. These neural measures include envelope reconstruction, integration window analysis, and TRF analysis. The results were aligned with previous findings that aging is associated with exaggerated cortical representations of speech (Decruy et al. 2019; Presacco et al. 2016a) and further investigated the cortical processing stages associated with this exaggeration. Using the integration window analysis and TRF peaks, we have now shown that all major cortical processing stages, early, middle and late processing, contributed to exaggerated neural responses. As previously shown, the addition of a competing talker diminishes the cortical representation of the attended speech signal, and here we have now also shown that aging enhances this reduction. In particular, TRF peak analysis has now revealed that it was only the middle and late processing contributions to the cortical representation that differ in amplitude between attended and unattended speech, and that difference was affected by age and the interfering speech in a complex manner. Additionally, TRF peak latency analysis has now shown that all processing stages were delayed with interfering speech, which was further affected by aging. Details of these novel findings are addressed below.

### Aging is associated with exaggerated speech envelope representation/encoding

Perhaps counterintuitively, the reconstruction analysis demonstrated that compared to younger adults, older adults exhibit exaggerated speech envelope representation irrespective of the noise condition. This replicates previous results by Presacco et al. (2016b) showing that older adults have a more robust (overly large) representation of the attended speech envelope in the cortex, and is consistent with studies showing enhanced envelope tracking with advancing age, both for discrete stimuli (Bidelman et al. 2014; Goossens et al. 2016; Irsik et al. 2021) and continuous speech stimuli (Decruy et al. 2018; Mesik et al. 2021). Whereas both Presacco et al. (2016b) and Decruy et al. (2018) analyzed the attended speech envelope reconstruction, the current study extends the analysis to the unattended speech envelope reconstruction. Incorporating both attended and unattended talker representations allows investigation of the two speech streams as distinct auditory objects (Griffiths and Warren 2004), separable via neural implementations of auditory scene analysis (Shinn-Cunningham 2008; Shamma et al. 2011). Here we have demonstrated that even the unattended speech envelope is exaggerated in the cortex of older adults and cortical exaggeration is not limited only to the attended speech. Moreover, the dynamical difference comparisons between age groups show that age-related exaggeration begins at latencies as early as 50–100 ms, and continues as late as 350 ms. This suggests that neural mechanisms underlying the exaggerated representation are active even in the earliest cortical stages, and some persist throughout the late processing stages.

The exaggerated envelope representation in older adults manifests as exaggerated TRF peak amplitudes at all three processing stages, M50_TRF_, M100_TRF_ and M200_TRF_, and for both attended and unattended speech. The enhanced M50_TRF_ in older adults also agrees with the integration window results above, that the exaggerated representation starts even at early cortical processing stages. Larger early cortical peaks (∼50 ms latency) in older adults have been reported using both EEG (McCullagh and Shinn 2013; Roque et al. 2019b) and MEG (Brodbeck et al. 2018; Zan et al. 2020), for both speech in quiet and in noisy conditions. Alain et al. (2014) suggested that this increased neural activity may be caused by excitatory/inhibitory imbalance, which is further in agreement with animal studies (McCullagh and Shinn 2013), and is consistent with other studies (Brodbeck et al. 2018). Larger cortical peaks at ∼100 ms latency (e.g., the M100_TRF_) in older adults have been reported using both EEG (McCullagh and Shinn 2013) and MEG (Zan et al. 2020), and the exaggeration is bigger for the attended speech compared to unattended. The exaggerated response at this middle processing stage has been associated with increased task-related effort (Rao et al. 2010). However, previous studies have reported that the M100 is enhanced in older adults for both active and passive listening paradigms, and also for simple stimuli (Tremblay et al. 2003; Sörös et al. 2009; Herrmann et al. 2018, 2022). This suggests that not just cognition per se but also other age-related functional changes may contribute to this enhancement, as elaborated in the next section. For longer latency cortical peaks (∼200 ms latency, e.g., the M200_TRF_), previous studies have reported an enhanced late peak in both EEG (O’Sullivan et al. 2015; Fiedler et al. 2019) and MEG (Zan et al. 2020) when the stimulus was continuous speech. No age-related enhancement was seen for this stage, however, for a gap-in-noise detection task (Alain and Snyder 2008; Lister et al. 2011). This may indicate that the late processing stage entails an additional stage of processing during speech comprehension that is not activated during simpler tasks such as tone processing.

### Possible mechanisms underlying exaggerated speech representation

Several potential mechanisms have been put forward to explain such exaggerated neural responses in older adults; not all of them necessarily apply for each of the three (early, middle and late) processing stages, for both attended and unattended talkers. One well-supported explanation is an imbalance between excitatory and inhibitory currents, where a reduction in inhibition would result in greater neural currents and their electromagnetic fields, but, due to the importance of inhibition for neuronal tuning, likely worse sensitivity, both temporally and spectrally (Herrmann and Butler 2021). Gamma-aminobutyric acid (GABA)-mediated inhibition plays a major role in maintaining synchrony and spectral sensitivity in auditory circuits. Both animal (Willott et al. 1991; Caspary et al. 1995; Hughes et al. 2010b; Caspary et al. 2013; Richardson et al. 2013; Parthasarathy et al. 2019; Ramamurthy and Recanzone 2020) and human (Lalwani et al. 2019; Ross et al. 2020; Dobri and Ross 2021; Harris et al. 2022) studies have reported a reduction in age-related inhibitory circuits and function. This mechanism could apply to any of the three main processing stages and for both attended and unattended talkers. Another possible contributor to the age-related amplitude increase is additional auditory processing due to age-related reduction in cortical connectivity: Peelle et al. (2010) found reduced coherence among cortical regions necessary to support speech comprehension, thus requiring multiple cortical regions to redundantly neurally process the same stimulus information, which would also result in increased extracranial neural responses. This top-down effect might be especially important for the middle and later processing stages. Finally, additional processing of the attended speech might arise from explicitly top-down compensatory mechanisms, where additional cortical regions would be recruited to support the early processing deficits (Wild et al. 2012; Pichora-Fuller et al. 2016; Rumschlag et al. 2022). Imaging studies have shown that older adults, even in the absence of elevated hearing thresholds, engage more extensive and diffuse regions of frontal cortex at relatively lower task loads than younger adults (for a review, see Kuchinsky and Vaden (2020)). Linking age-related changes in neural activity to listening performance is critical for understanding the extent to which observed upregulation of activity is evidence of a compensatory process (generally predictive of better listening performance) or of an inability to inhibit irrelevant cortical processing (i.e., de-differentiation, predictive of poor performance; for a review, see Wingfield and Grossman (2006))

### Selective attentional modulation of speech representation/encoding and aging

In line with the results for younger adults (Mesgarani and Chang 2012; Ding and Simon 2012; O’Sullivan et al. 2015; Das, Bertrand, and Francart 2018), we have found that in older adults the attended speech envelope is better represented than the unattended. Surprisingly, irrespective of exaggerated envelope representation in the cortex, we have shown that both groups showed similar effects of selective attention on the envelope representation as no age by selective attention interaction effect was found.

Both integration window analysis and TRF analysis contrasting attended vs unattended speech have here revealed that the unattended speech is represented to a similar degree as the attended (or even more strongly in the case of −6 dB SNR, when it is acoustically louder), at the early processing stage. However, by the middle and late processing stages, attended speech is better represented than unattended in both groups. Therefore, the early processing stages better reflect the full acoustic sound scene than the selective-attention-driven percept, whereas the middle and later stages more closely follow the percept (though see Brodbeck et al. 2020). Older adults exhibit an enhanced attended-unattended M100_TRF_ amplitude difference compared to younger adults (in addition to showing enhancement in both separately), which may reflect task-related increased attention or cognitive effort that further supports the selective attention.

### Representations of quiet and noisy speech are differentially affected by age

Our analysis also investigated how competing speakers affect the cortical speech representation. We found that, irrespective of age, masking by other speakers adversely affects the cortical representation of the attended speech envelope. This is also in line with the previous studies, which report that envelope tracking is adversely affected by the SNR (Ding and Simon 2013; Presacco et al. 2016a,b; Das et al. 2018). Our results additionally show that older adults’ envelope representation is more strongly affected, still negatively, by adding higher levels of noise, compared to younger adults. Thus, within an individual, better reconstruction accuracy is associated with clearer speech, but not across age groups; for example, older adults’ reconstruction accuracy for 0 dB SNR is typically better than younger adults’ reconstruction accuracy for quiet (as seen in Figure 3(a)).

A drop in M50_TRF_ amplitude from quiet to noisy speech, and with similar TRF peak amplitudes for both attended and unattended speech at 0 dB SNR, is expected for an early auditory cortical stage that processes the complete acoustic scene and thus both attended and unattended talkers (Fiedler et al. 2019; Brodbeck et al. 2020). The older adults here showed greater reduction in M50 _TRF_ amplitude with the noise condition, suggesting that aging adversely affects the early-stage cortical processing in speech-in-noise conditions. In contrast, older adults display increasing M100_TRF_ amplitude with the noise condition, whereas no significant M100_TRF_ amplitude differences are seen between noise conditions in younger adults. These results are consistent with some previous studies (Ding and Simon 2012b; McCullagh et al. 2012; Rufener et al. 2014), but not all (McCullagh and Shinn 2013; Billings et al. 2015; Zan et al. 2020) where a dependence on masker noise condition is found in both groups. Mechanistically, an exaggerated M100_TRF_ amplitude may reflect an increase in task-related attention or cognitive effort (Rao et al. 2010; Billings et al. 2015), and as such any conflicting trends maybe due to task difference subtleties. In contrast, the M200_TRF_ amplitude, also has a pronounced decrease from quiet speech to noisy speech in older adults (no such drop is observed in younger adults). The M200_TRF_ amplitude is also strongly enhanced by selective attention, and it has a sufficiently long latency to reflect top-down compensatory processing known to be important for older adults (Pichora-Fuller 2008). One possibility for the noise-related decrease in older adults is that the observed M200_TRF_ actually reflects the sum of two sources with similar latencies but opposite polarities, where the first (positive) source is active regardless of whether the speech is noisy, but the second (negative and slightly later) top-down compensatory source is invoked only under difficult listening conditions; this finding is also consistent with the association between decreased M200_TRF_ peak amplitudes and longer latencies in older adults. This late peak in older adults has also been reported as a potential biomarker for *behavioral inhibition* (Zan et al. 2020), though the current study did not find any such correlations between M200_TRF_ amplitude and behavioral measures. The different amplitude trends as a function of masker level, between the M100_TRF_ and M200_TRF_ for the two groups, indicate that the presence and level of the masker significantly contributes to middle and late processing in older adults.

### Aging is associated with earlier early processing and prolonged late processing

The integration window analysis revealed that a long temporal integration window (at least ∼300 ms) allows a robust speech reconstruction for both groups, which is consistent with early studies using only younger adults (Ding and Simon 2013; O’Sullivan et al. 2015). Specifically, Power et al. (2012) and O’Sullivan et al. (2015) reported that an interval of duration 170 – 250 ms is important for attention decoding with EEG, and processing at that latency may even be at the level of semantic analysis. The striking difference between the age groups, however, is that older adults need more time to better represent the speech envelope, as seen in Figure 4(c), as long as 350 ms (Presacco et al. 2016a). This provides support for an existence of late compensatory mechanisms to support the additional selective attention filtering process, required for the early-stage processing deficits or slowing of synchronous neural firing rate (Tremblay et al. 2004; O’Brien et al. 2015), which is addressed explicitly in the TRF analysis discussed next.

TRF peak latencies indicate processing time needed to generate responses after the corresponding acoustic feature, and so can be mapped to the speed of auditory processing. Both age groups showed significant noise-related delays in the M50_TRF_, M100_TRF_ and M200_TRF_ peak latencies, suggesting longer cortical processing associated with the addition (and level) of the masker (McCullagh and Shinn 2013). The latency results for babble speech were less clear. Latencies for the babble condition were delayed compared to quiet speech, yet no consistent trends were observed compared to two-talker conditions. The babble speech may be impossibly challenging for some listeners who are more likely to disengage attention, reducing top-down effects, but for other listeners its challenge may not exceed limits, enhancing top-down effects, thus confounding comparisons between listeners (Kuchinsky and Vaden 2020).

Compared to younger adults, older adults demonstrated relatively early M50_TRF_ peaks. This effect has been observed in some studies using both CAEP (O’Brien et al. 2015; Roque et al. 2019b) and MEG (Brodbeck et al. 2018), but not others (Tremblay et al. 2004; Alain et al. 2014). This finding is consistent with an excitation/inhibition imbalance favoring excitation compared to younger adults. Another possible explanation is that an M50_TRF_ followed immediately by an exaggerated M100_TRF_ of the opposite polarity would appear shortened due to an earlier cut off imposed by the subsequent peak, with the artifactual side-effect of shorter latency. The M100_TRF_ latency was comparable for both groups, which contrasts with the late peak M200_TRF_ which was significantly delayed in older adults. These findings are in line with studies that recorded responses to speech syllables (Tremblay and Newman 2015; Roque et al. 2019a), suggesting that an age-related decrease in rate of transmission for auditory neurons contributing to P2. In contrast to the early peak, both middle and late peaks demonstrated further delayed peak latencies with noise condition in older adults suggesting over-reliance on the middle and late processing mechanisms in older adults to compensate for degraded afferent input (Parthasarathy et al. 2020). Taken together, age-related impaired processing of the auditory input at the early stages could affect the auditory scene representation at the late processing stages, by employing additional cortical regions and compensatory mechanisms at the later stage.

Our results also add additional supporting evidence that the attended speech signal requires longer processing times in later stages to discern the information in the attended stream in older listeners, possibly to recover the early processing deficits by many compensatory mechanisms. Fiedler et al. (2019), using EEG, demonstrated that late cortical tracking of the unattended talker reflects neural selectivity in acoustically challenging conditions. They reported an early suppressed positive P2 in line with the current study and additional late negative N2 peak for the unattended talker which appears around the same latency as attended P2. They argue that this late N2 of the unattended talker actively suppresses the distracting inputs.

### Behavior

As expected, SPIN scores and intelligibility ratings decreased as noise condition increased in both groups, suggesting that noise condition negatively affects speech intelligibility and in turn increases task difficulty. However, no significant difference was found between younger and older groups in the SPIN scores, which was unexpected. The SPIN measure employed here, developed from the same materials used during the MEG recordings, has not been calibrated against more standard SPIN measures and so may not be able to distinguish the hearing complications which arise with aging. For this task, subjects listened to a very short narrative segments (with 4–7 key words) with no time limit, and the older subjects may have benefited not only from their extended vocabulary and language experience (Pichora-Fuller et al. 1995; Pichora-Fuller 2008; Schneider et al. 2016) but also from the lack of time demands. In addition, the range of scores obtained in both groups suggests that the task was more challenging than other standardized measures, especially as some of the younger listeners did not achieve 100% performance even in the quiet condition. In contrast, established speech intelligibility tests such as the QuickSIN (Killion et al. 2004) are known to show such behavioral age-related auditory declines (Presacco et al. 2016a,b; Holder et al. 2018). Unfortunately, due to the COVID-19 pandemic, QuickSIN measures were obtained from only half of the subjects, which were not enough to incorporate into analysis. The subjective intelligibility ratings of older adults were, perhaps surprisingly, higher than those of younger adults. This finding is nevertheless consistent with earlier results showing that the older adults tend to underestimate their hearing difficulties relative to younger adults (Uchida et al. 2003; Gosselin and Gagné 2011). However, it is unclear whether it is younger adults that overestimate, or that older adults underestimate, their hearing difficulties (or both), and whether their subjective judgements are differently influenced by intelligibility, contextual factors, or other variables (Perron et al. 2022).

The positive association between the behavioral SPIN scores and intelligibility ratings do suggest that models using (subjective) intelligibility ratings collected during the task of interest would give similar outcomes as those using (objective) SPIN scores collected in a separate task (Hazan et al. 2018). However, the age effects revealed only in subjective intelligibility ratings suggests that the two measures reflect different aspects of speech intelligibility or different factors that affect performance.

Our results also revealed that within a subject, the neural measures of reconstruction accuracy M50_TRF,_ and TRF peak amplitude are correlated with speech intelligibility (though speech intelligibility cannot be disentangled here from the changes in noise condition). The positive association between intelligibility and larger neural measures in general is suggestive of a compensatory (vs. de-differentiation) pattern of neural engagement, in which additional resources are brough online to maintain performance in challenging conditions (Wingfield and Grossman 2006). Age-related compensation may be particularly evident during later-stage processing, as the correlation between the M200_TRF_ peak amplitude and better speech intelligibility score was only observed for older adults. Unexpectedly, we did not find any consistent relationships between neural measures and behavioral performance measures within a noise condition. One possible reason, as mentioned above, is that these uncalibrated behavioral performance measures may not sufficiently capture the known age-related hearing difficulties and temporal processing deficits.

### Audiometry Differences

In the current study, we investigated the age related cortical temporal processing deficits by recruiting younger and older participants who have clinically normal hearing (pure-tone thresholds ≤ 25 dB hearing level (HL) from 125 to 4000 Hz). However, it is difficult to completely eliminate the confounds of peripheral hearing loss that gradually occur with aging. Previous studies have reported that this peripheral hearing loss may contribute to some of the speech-in-noise understanding difficulties experienced by older adults (Motlagh Zadeh et al. 2019; Yeend et al. 2019). The average of the pure-tone thresholds (PTA) showed that there is a positive correlation between the age and PTA (Pearson’s *r* = 0.87, *p* < 0.001). Therefore, the causal relationship between the hearing sensitivity and the aging may cast doubt on the age-related neural changes we observed in the current study. To answer this issue, we investigated the effects of hearing sensitivity (PTA) on the neural and behavioral measures analyzed in the current study. As there is a significant correlation between age and PTA, the LMER models wipes out the PTA from the models. Therefore, we evaluated the effects of PTA within an age group separately. However, we did not find any significant correlation between PTA and neural measures (reconstruction accuracy, TRF amplitudes and latencies) nor PTA and behavioral measures (intelligibility ratings and SPIN scores).

Age-related deterioration both peripherally, such as subclinical loss of outer and inner hair cells and ganglion cells within the cochlea (Parthasarathy and Kujawa 2018; Wu et al. 2019), and centrally, such as loss of neural synchrony (Boettcher et al. 1993; Schneider and Pichora-Fuller 2000; Chisolm et al. 2003; Anderson and Karawani 2020) may not affect audiometric thresholds, but likely contribute to suprathreshold listening difficulties (Plack et al. 2014). Previous studies including both hearing impaired and normal hearing older adults have shown that beyond hearing sensitivity, aging is the main driving factor of temporal processing differences in subcortical and cortical regions (Presacco et al. 2019; Roque et al. 2019a). Therefore, these results suggest that, aging and age-related hearing difficulties other than hearing sensitivity may cause the observed temporal processing differences in the auditory cortex. However, we acknowledge that audiometric thresholds may not adequately characterize peripheral auditory function, and that the inclusion of other measures of peripheral function in the analysis, such as otoacoustic emissions or auditory brainstem response Wave I amplitude, may reveal central consequences of decreased afferent input (Anderson et al. 2021).

## CONCLUSION

In conclusion, the present study showed that aging is associated with exaggerated speech representation in the cortical response and this exaggeration is noted in all three cortical processing stages; early, middle and late processing. Moreover, the effects of speech intelligibility and attention on the respective three M50_TRF_, M100_TRF_ and M200_TRF_ peak amplitudes and latencies reveal characteristics related to different cortical processing stages, and aging appears to differently affect the individual processing stages. Earlier and enhanced processing of early stages support the hypothesis of an excitatory and inhibitory imbalance in older adults. The underlying causes of enhanced processing of the middle cortical processing stage are less clear; indeed, the strongest enhancement occurs under noise conditions where the timing is neither earlier nor later than for younger adults. The late auditory cortical stage displays both delayed and enhanced processing in older adults, which were associated with better performance on a parallel speech-in-noise task and thus consistent with cognitive (perhaps memory/experience based) compensatory mechanisms. Overall, these findings support the theory that that some of the age-related difficulties in understanding speech in noise experienced by older adults, including those directly related to temporal processing, are accompanied by age-related temporal processing differences in auditory cortex.

## ACKNOWLEDGEMENTS

This work was supported by grants from the National Institute on Aging (P01-AG055365) and the; National Institutes of Deafness and Other Communication Disorders (R01-DC014085, R01-DC019394). We thank Katie Brow, Julie Cohen and Carol Gorham for their help in recruiting and testing subjects.

## AUTHOR CONTRIBUTIONS

I.M.D.K., J.L.D, J.P., A.P., S.A., S.E.K., and J.Z.S. conceived and designed the research; I.M.D.K., J.L.D, J.P., and L.D. performed experiments; I.M.D.K., J.L.D., L.D. and J.Z.S. analyzed data; I.M.D.K., L.D., S.A., S.E.K., and J.Z.S. interpreted results of experiments; I.M.D.K. prepared figures; I.M.D.K. drafted manuscript; I.M.D.K., J.L.D, J.P., A.P., L. D., S.A., S.E.K., and J.Z.S. edited and revised manuscript; I.M.D.K., J.L.D, J.P., A.P., L. D., S.A., S.E.K., and J.Z.S. approved final version of manuscript.

## APPENDIX

**Table A1.** LMEM summary tables for behavioral analysis

|  | SPIN score |  |  |  | Intelligibility Rating |  |  |  |
| --- | --- | --- | --- | --- | --- | --- | --- | --- |
| *Fixed Effects* | *Estimates* | *SE* | *t value* | *p value* | *Estimates* | *SE* | *t value* | *p value* |
| Intercept [Younger, Quiet] | 93.59 | 1.92 | 48.6 | <0.001 | 8.72 | 0.27 | 32.2 | <0.001 |
| noise cond. [0 dB] | -12.88 | 1.57 | -8.2 | <0.001 | -2.60 | 0.27 | -9.5 | <0.001 |
| noise cond. [-6 dB] | -53.83 | 1.57 | -34.3 | <0.001 | -4.10 | 0.34 | -12.0 | <0.001 |
| noise cond. [Bab] | -67.35 | 1.92 | -35.1 | <0.001 | -4.63 | 0.43 | -10.9 | <0.001 |
| age [Older] |  |  |  |  | 0.76 | 0.36 | 2.1 | 0.035 |
| *Random Effects* | *Variance* | *SD* |  |  | *Variance* | *SD* |  |  |
| Intercept \| subject | 76.6 | 8.76 |  |  | 0.88 | 0.94 |  |  |
| noise cond. [0 dB] \| subject |  |  |  |  | 1.29 | 1.14 |  |  |
| noise cond. [-6 dB] \| subject |  |  |  |  | 2.72 | 1.65 |  |  |
| noise cond. [Bab] \| subject |  |  |  |  | 4.32 | 2.08 |  |  |
| Number of obs.: 217, Subjects: 31 |  |  |  |  | Number of obs.: 238, Subjects: 34 |  |  |  |

*SPIN score* ~ 1 + noise cond. + (1|subject)
*Intelligibility Rating* ~ 1 + noise cond. + age + (1 + noise cond. |subject)

**Table A2.** LMEM summary table for SPIN score vs Intelligibility score

| SPIN score |  |  |  |  |
| --- | --- | --- | --- | --- |
| *Fixed Effects* | *Estimates* | *SE* | *t value* | *p value* |
| (Intercept) | 12.08 | 4.22 | 2.9 | 0.005 |
| Intelligibility rating | 8.2 | 0.58 | 14.2 | <0.001 |
| Random Effects |  |  | Variance | SD |
| Intercept \| subject | 71.98 |  | 8.49 |  |
| Number of obs.: 217, Subjects: 31 |  |  |  |  |

*SPIN score* ~ Intelligibility rating + (1|subject)

**Table A3.**
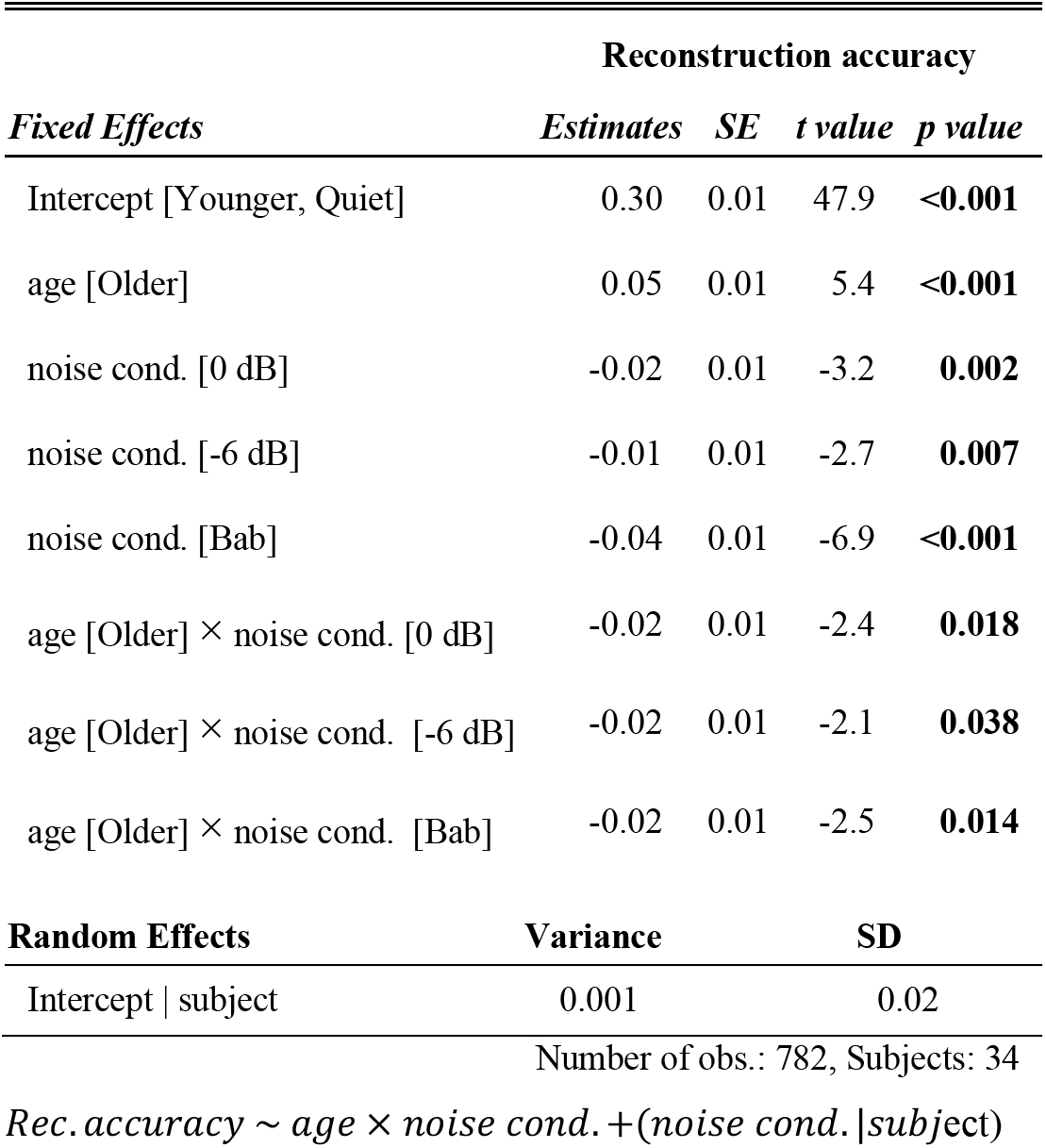
LMEM summary table for attended speech envelope reconstruction accuracies

**Table A4.** LMEM summary table for attended vs unattended speech envelope reconstruction accuracies

|  | Reconstruction accuracy |  |  |  |
| --- | --- | --- | --- | --- |
| *Fixed Effects* | *Estimates* | *SE* | *t value* | *p value* |
| Intercept [Attended, Younger] | 0.29 | 0.01 | 49.9 | <0.001 |
| attention [Unattended] | -0.03 | 0.00 | -6.9 | <0.001 |
| age [Older] | 0.03 | 0.01 | 4.1 | <0.001 |
| Random Effects | Variance | SD |  |  |
| Intercept \| subject | 0.0007 | 0.03 |  |  |
| attention \| subject | 0.0003 | 0.02 |  |  |
| Number of obs.: 816, Subjects: 34 |  |  |  |  |
| *Rec.accuracy* ~ *age* + *attention* + (1 + *attention*\|*subject*) |  |  |  |  |

**Table A5.** LMEM summary table for attended TRF peak amplitudes

|  | M50<sub>TRF</sub> amplitude |  |  |  | M100<sub>TRF</sub> amplitude |  |  |  | M200<sub>TRF</sub> amplitude |  |  |  |
| --- | --- | --- | --- | --- | --- | --- | --- | --- | --- | --- | --- | --- |
| *Fixed Effects* | *Est.* | *SE* | *t value* | *p value* | *Est.* | *SE* | *t value* | *p value* | *Est.* | *SE* | *t value* | *p value* |
| Intercept<br>[Younger, Quiet] | 0.02 | 0.003 | 5.6 | <0.001 | 0.02 | 0.004 | 5.2 | <0.001 | 0.01 | 0.003 | 2.3 | 0.021 |
| age [Older] | 0.02 | 0.01 | 3.5 | <0.001 | 0.01 | 0.01 | 0.9 | 0.369 | 0.03 | 0.004 | 6.8 | <0.001 |
| noise cond.[0 dB] | -0.01 | 0.003 | -1.6 | 0.101 | -0.00 | 0.003 | -0.6 | 0.567 | -0.00 | 0.003 | -0.1 | 0.931 |
| noise cond.[-6 dB] | -0.01 | 0.003 | -2.1 | 0.038 | -0.00 | 0.003 | -0.3 | 0.745 | -0.00 | 0.003 | -0.5 | 0.612 |
| noise cond.[Bab] | -0.02 | 0.003 | -5.2 | <0.001 | -0.01 | 0.003 | -2.6 | 0.008 | -0.00 | 0.003 | -1.1 | 0.289 |
| age [Older] ×<br>noise cond.[0 dB] | -0.01 | 0.005 | -2.1 | 0.034 | 0.01 | 0.005 | 2.9 | 0.004 | -0.02 | 0.004 | -4.8 | <0.001 |
| age [Older] ×<br>noise cond.[-6 dB] | -0.01 | 0.005 | -2.9 | 0.004 | 0.01 | 0.005 | 2.6 | 0.009 | -0.02 | 0.004 | -4.9 | <0.001 |
| age [Older] ×<br>noise cond.[Bab] | -0.01 | 0.005 | -2.2 | 0.029 | 0.01 | 0.005 | 2.1 | 0.036 | -0.03 | 0.004 | -5.8 | <0.001 |
| **Random Effects** | **Variance** |  | **SD** |  | **Variance** |  | **SD** |  | **Variance** |  | **SD** |  |
| Intercept \| subject | 0.0001 |  | 0.01 |  | 0.0002 |  | 0.01 |  | 0.0001 |  | 0.01 |  |
| Number of obs.: 136, Subjects: 34 |  |  |  | Number of obs.: 136, Subjects: 34 |  |  |  | Number of obs.: 136, Subjects: 34 |  |  |  |  |
| $M50_{TRF} \sim age \times noise\ cond. + (1\vert subject)$ | | | | | | | | | | | | |
| $M100_{TRF} \sim age \times noise\ cond. + (1\vert subject)$ | | | | | | | | | | | | |
| $M200_{TRF} \sim age \times noise\ cond. + (1\vert subject)$ | | | | | | | | | | | | |

**Table A6.** LMEM summary table for attended vs unattended TRF peak amplitudes

| *Fixed Effects* | **M50<sub>TRF</sub> amplitude** |  |  |  | **M100<sub>TRF</sub> amplitude** |  |  |  | **M200<sub>TRF</sub> amplitude** |  |  |  |
| --- | --- | --- | --- | --- | --- | --- | --- | --- | --- | --- | --- | --- |
|  | *Est.* | *SE* | *t value* | *p value* | *Est.* | *SE* | *t value* | *p value* | *Est.* | *SE* | *t value* | *p value* |
| Intercept<br>[Attended,<br>Younger, 0 dB] | 0.02 | 0.000 | 7.6 | <0.001 | 0.02 | 0.000 | 6.1 | <0.001 | 0.01 | 0.000 | 4.8 | <0.001 |
| attention<br>[Unattended] | 0.01 | 0.00 | 5.8 | <0.001 | -0.01 | 0.00 | -5.9 | <0.001 | -0.01 | 0.00 | -4.7 | <0.001 |
| age [Older] |  |  |  |  | 0.02 | 0.00 | 3.8 | <0.001 | 0.01 | 0.00 | 3.7 | <0.001 |
| attention<br>[Unattended] ×<br>age [Older] |  |  |  |  | -0.01 | 0.00 | -4.1 | <0.001 |  |  |  |  |
| noise cond.[-6 dB] |  |  |  |  |  |  |  |  | -0.00 | 0.00 | -1.0 | 0.310 |
| attention<br>[Unattended] ×<br>noise cond.[-6 dB] |  |  |  |  |  |  |  |  | 0.00 | 0.00 | 1.2 | 0.237 |
| **Random Effects** | **Variance** |  | **SD** |  | **Variance** |  | **SD** |  | **Variance** |  | **SD** |  |
| Intercept \| subject | 0.0001 |  | 0.01 |  | 0.0001 |  | 0.01 |  | 0.0001 |  | 0.00 |  |

Number of obs.: 136, Subjects: 34 Number of obs.: 136, Subjects: 34 Number of obs.: 136, Subjects: 34
$M50_{TRF} \sim attention + (1|subject)$
$M100_{TRF} \sim age \times attention + (1|subject)$
$M200_{TRF} \sim age \times noise\ cond. + (1|subject)$

**Table A7.** LMEM summary table for attended TRF peak latencies

|  | M50<sub>TRF</sub> Latency |  |  |  | M100<sub>TRF</sub> Latency |  |  |  | M200<sub>TRF</sub> Latency |  |  |  |
| --- | --- | --- | --- | --- | --- | --- | --- | --- | --- | --- | --- | --- |
| *Fixed Effects* | *Est.* | *SE* | *t value* | *p value* | *Est.* | *SE* | *t value* | *p value* | *Est.* | *SE* | *t value* | *p value* |
| Intercept<br>[Younger, Quiet] | 42.44 | 2.85 | 14.9 | <0.001 | 120.57 | 3.32 | 36.3 | <0.001 | 178.70 | 5.20 | 34.4 | <0.001 |
| age [Older] | -2.27 | 4.11 | -0.6 | 0.581 | -9.44 | 4.84 | -1.9 | 0.051 | 11.05 | 7.22 | 1.5 | 0.126 |
| noise cond.[0 dB] | 18.48 | 3.02 | 6.1 | <0.001 | 14.59 | 2.75 | 5.3 | <0.001 | 21.80 | 4.64 | 4.7 | <0.001 |
| noise cond.[-6 dB] | 13.07 | 2.99 | 4.4 | <0.001 | 17.65 | 2.75 | 6.4 | <0.001 | 29.97 | 4.91 | 6.1 | <0.001 |
| noise cond.[Bab] | 25.63 | 3.12 | 8.2 | <0.001 | 16.99 | 2.73 | 6.2 | <0.001 | 30.82 | 5.12 | 6.0 | <0.001 |
| noise cond.[0 dB]<br>× age [Older] | -7.66 | 4.32 | -1.8 | 0.076 | 8.05 | 3.98 | 2.0 | 0.043 | 17.08 | 6.68 | 2.6 | 0.011 |
| noise cond.[-6 dB]<br>× age [Older] | -0.49 | 4.30 | -0.1 | 0.909 | 11.23 | 3.97 | 2.8 | 0.004 | 11.16 | 6.97 | 1.6 | 0.109 |
| noise cond.[Bab] ×<br>age [Older] | -16.39 | 4.54 | -3.6 | <0.001 | 8.64 | 3.95 | 2.2 | 0.023 | -6.92 | 7.11 | -0.9 | 0.330 |
| Random Effects | Variance |  | SD |  | Variance |  | SD |  | Variance |  | SD |  |
| Intercept \| subject | 60.9 |  | 7.8 |  | 128.2 |  | 11.32 |  | 270.8. |  | 16.46 |  |
| Number of obs.: 125, Subjects: 34 Number of obs.: 131, Subjects: 34 Number of obs.: 109, Subjects: 32 |  |  |  |  |  |  |  |  |  |  |  |  |
| $M50_{TRF} \sim age \times noise\ cond. + (1\vert subject)$ | | | | | | | | | | | | |
| $M100_{TRF} \sim age \times noise\ cond. + (1\vert subject)$ | | | | | | | | | | | | |
| $M200_{TRF} \sim age \times noise\ cond. + (1\vert subject)$ | | | | | | | | | | | | |

**Table A8.** LMEM summary table for attended vs unattended TRF peak latencies

| *Fixed Effects* | **M50<sub>TRF</sub> Latency** |  |  |  | **M100<sub>TRF</sub> Latency** |  |  |  | **M200<sub>TRF</sub> Latency** |  |  |  |
| --- | --- | --- | --- | --- | --- | --- | --- | --- | --- | --- | --- | --- |
|  | *Estimates* | *SE* | *t value* | *p value* | *Estimates* | *SE* | *t value* | *p value* | *Estimates* | *SE* | *t value* | *p value* |
| Intercept [Younger, Attended] | 60.76 | 2.79 | 21.8 | **<0.001** | 136.62 | 2.73 | 50.1 | **<0.001** | 202.15 | 4.90 | 41.3 | **<0.001** |
| age [Older] | -9.76 | 4.03 | -2.4 | **0.015** | -1.97 | 3.93 | -0.5 | 0.616 | 27.84 | 7.09 | 3.9 | **<0.001** |
| attention [Unattended] | 0.50 | 3.11 | 0.2 | 0.873 | 8.03 | 3.21 | 2.5 | **0.012** | -12.02 | 6.28 | -1.9 | 0.056 |
| noise cond.[-6 dB] | -5.40 | 2.17 | -2.5 | **0.013** | 2.95 | 1.46 | 2.0 | **0.043** | 7.15 | 2.38 | 3.0 | **0.003** |
| age [Older] × attention [Unattended] | -2.25 | 4.48 | -0.5 | 0.616 | -12.01 | 4.56 | -2.6 | **0.008** | -28.54 | 8.26 | -3.5 | **0.001** |
| attention [Unattended] × noise cond.[-6 dB] | 6.80 | 3.03 | 2.3 | **0.025** |  |  |  |  | -12.04 | 3.96 | -3.0 | **0.002** |
| age [Older] × noise cond.[-6 dB] | 7.15 | 3.08 | 2.3 | **0.020** |  |  |  |  |  |  |  |  |
| age [Older] × attention [Unattended] × noise cond.[-6 dB] | -9.05 | 4.33 | -2.1 | **0.037** |  |  |  |  |  |  |  |  |
| **Random Effects** | **Variance** |  | **SD** |  | **Variance** |  | **SD** |  | **Variance** |  | **SD** |  |
| Intercept \| subject | 97.58 |  | 9.88 |  | 101.24 |  | 10.06 |  | 330.42 |  | 18.18 |  |
| attention [Unattended] \| subject | 89.56 |  | 9.43 |  | 117.18 |  | 10.82 |  | 313.69 |  | 17.71 |  |
| noise cond.[-6 dB] \| subject |  |  |  |  | 31.28 |  | 5.59 |  |  |  |  |  |

Number of obs.: 133, Subjects: 34 Number of obs.: 116, Subjects: 34 Number of obs.: 94, Subjects: 32
$M50_{TRF} \sim age \times attention \times noise\ cond. + (1 + attention|Subject)$
$M100_{TRF} \sim age \times attention + noise\ cond. + (1 + attention + noise\ level|subject)$
$M200_{TRF} \sim age \times attention + attention \times noise\ cond. + (1 + attention|subject)$

**Table A9.** LM summary table for M200_TRF_ amplitudes vs latency

|  | 0 dB |  |  |  | -6 dB |  |  |  |
| --- | --- | --- | --- | --- | --- | --- | --- | --- |
| *Effects* | *Estimates* | *SE* | *t value* | *p value* | *Estimates* | *SE* | *t value* | *p value* |
| Intercept [Younger] | -0.04 | 0.03 | -1.4 | 0.17 | -0.06 | 0.04 | -1.5 | 0.14 |
| age [Oder] | 0.08 | 0.03 | 2.8 | <0.001 | 0.13 | 0.05 | 2.9 | 0.01 |
| latency | 0.0002 | 0.0001 | 1.7 | 0.1 | 0.0003 | 0.0002 | 1.7 | 0.13 |
| age [Older] × latency | -0.0004 | 0.0001 | -2.4 | 0.025 | -0.0006 | 0.0002 | -2.6 | 0.015 |
| Subjects: 30 |  |  |  | Subjects: 26 |  |  |  |  |

$M200_{TRF}amplitude \sim age \times M200_{TRF}latency$

**Table A10.** GAMM summary table for Integration Window Analysis

| Parametric Coefficients | Reconstruction Accuracy |  |  |  |
| --- | --- | --- | --- | --- |
|  | *Estimates* | *SE* | *t value* | *p value* |
| Intercept [Younger, Quiet] | 0.25 | 0.00 | 55.42 | <0.001 |
| Older × Quiet | 0.04 | 0.01 | 6.43 | <0.001 |
| Younger × 0 dB × Attended | -0.02 | 0.001 | -4.56 | <0.001 |
| Older × 0 dB × Attended | 0.01 | 0.01 | 1.45 | 0.146 |
| Younger × 0 dB × Unattended | -0.03 | 0.001 | -8.21 | <0.001 |
| Older × 0 dB × Unattended | -0.01 | 0.01 | -1.78 | 0.075 |
| Younger × -6 dB × Attended | -0.01 | 0.001 | -3.50 | <0.001 |
| Older × -6 dB × Attended | 0.02 | 0.01 | 2.29 | 0.022 |
| Younger × -6 dB × Unattended | -0.03 | 0.001 | -8.19 | <0.001 |
| Older × -6 dB × Unattended | -0.00 | 0.01 | -0.11 | 0.914 |
| Smooth Terms | edf |  | Ref.df | *p* |
| s(Wind) : Younger × Quiet | 5.93 | | 370.86 | <0.001 |
| s(Wind) : Older × Quiet | 5.94 | | 520.27 | <0.001 |
| s(Wind) : Younger × 0 dB × Attended | 5.80 | | 243.32 | <0.001 |
| s(Wind) : Older × 0 dB × Attended | 5.90 | | 336.61 | <0.001 |
| s(Wind) : Younger × 0 dB × Unattended | 5.89 | | 231.45 | <0.001 |
| s(Wind) : Older × 0 dB × Unattended | 5.94 | | 298.78 | <0.001 |
| s(Wind) : Younger × -6 dB × Attended | 5.82 | | 249.48 | <0.001 |
| s(Wind) : Older × -6 dB × Attended | 5.86 | | 330.57 | <0.001 |
| s(Wind) : Younger × -6 dB × Unattended | 5.91 | | 227.02 | <0.001 |
| s(Wind) : Older × -6 dB × Unattended | 5.94 | | 302.23 | <0.001 |
| s (Wind, Subject) | 107.49 |  | 5.68 | <0.001 |

Number of obs.: 14144, Subjects: 34

